# The “five divergent amino acids rule” and evolution analysis of cytochrome *c* oxidase as new methods for *Scolopendra* Linnaeus, 1758 species differentiation (Chilopoda. Scolopendromorpha)

**DOI:** 10.1101/2025.11.24.690109

**Authors:** Carles Doménech, Víctor M. Barberá, Eduardo Larriba

## Abstract

*Scolopendra* Linnaeus, 1758 is a genus of centipede showing reasonably good morpho-molecular correlation. This often allows the acceptable species differentiation and phylogenetic positioning by just using the partial sequence of the cytochrome *c* oxidase (COI) gene. However, for biological, statistical, or technical reasons, several exceptions to that fact have been observed, which together with a lack of a solid criterion to differentiate species at the molecular level, make difficult the interpretation of some taxonomic and systematic outcomes.

With the scope of providing a standardized system improving the molecular *Scolopendra* species delimitation and surpassing some issues related with the mtDNA’s use, a total of 45 representative COI sequences belonging to 22 taxa are tested, but for the first time, performing distinct amino acid’s chains evolution analyses. To illustrate this, a general evaluation of the genus is firstly provided, while to exemplify some of the deeper analysis, the cases of *S. paradoxa* Doménech, 2018 and *S. spinosissima* Kraepelin, 1903 are then here re-explored using some alternative tools.

As a result, the partial COI protein sequences analysis identified, at least, ten amino acid residues’ positions as useful for *Scolopendra* species differentiation, being generally five divergent residues enough to distinguish taxa [“the five divergent amino acid rule”]. When this premise wasn’t fulfilled, the amino acid frequency, exclusivity, or electrochemical properties and especially the *r* substitution probability test, helped solve the cases. Among *S. paradoxa* and *S. spinosissima* five diverging residues were found, with one of them being exclusive to the entire genus for the former taxon. Also, the substitution probability (*r*) analysis showed strong positive selection for three of these five divergent amino acids.

As exceptions, *S. dawydoffi* Kronmüller, 2012 has been found indistinguishable from *S. multidens* Newport, 1844 by this method, suggesting its eventual synonymy, while *S. cataracta* Siriwut, Edgecombe & Panha, 2016 showed an intraspecific maximum divergence of 5 amino acids. Finally and at nomenclatural level, the species name *S. hainanum* Kronmüller, 2012 is emended as *S. hainanensis* while the taxon *S. mojiangica* Zhang & Chi, is here declared *nomen nudum*, due its original description remains unavailable.

## Introduction

The class Chilopoda is among the oldest groups of terrestrial arthropods (Edgecombe & Giribet 2007). These soil dweller invertebrates’ (Undheim & King 2011; Undheim et al. 2015) are venomous predators widely distributed across the warmer and tropical ecosystems. They are classified into five distinct orders, which currently include a total of 3,300 described taxa (Edgecombe & Giribet 2007). Among the 700 valid species of the order Scolopendromorpha, the most recognizable genus is *Scolopendra* Linnaeus, 1758, which currently comprises around 100 accepted members (Bonato et al. 2016; Schileyko et al. 2020).

Like many other metazoans and since their first descriptions, species of this genus have undergone multiple taxonomic and nomenclatural changes (see, for example, Linnaeus 1758; Kraepelin 1903; Attems 1930; Kronmüller 2012). However, these classical classification systems, based mainly on morphological and geographical differences, have been significantly improved by integrating them with more modern molecular techniques. As a paradigm of this progress, cytochrome *c* oxidase (COI) has proven to be a reliable bioindicator capable of establishing consistent morpho-molecular phylogenies across various groups. These have helped resolve some of the difficult relationships within the genus *Scolopendra* (Edgecombe & Giribet 2008; Vahtera et al. 2012, 2013; Oeyen et al. 2014; Siriwut et al. 2015). Likewise, through the quantitative analysis of genetic distances in partial COI sequences, it has also been possible to distinguish species with a reasonable degree of confidence (Siriwut et al. 2016; Doménech et al. 2018). However, despite generally working well, these methods do not always reflect an expected correlation between the morphology of all taxa and their percentage of divergence or phylogenetic position. For example, some studies have revealed that two congeners, despite being morphologically very different, appear to be more molecularly similar to each other than to their morphologically closer relatives. In some cases, even species placed in different genera turned out to be more similar to each other at the molecular level than to their actual congeners (Vahtera et al. 2012, 2013; Vahtera & Edgecombe 2014; Siriwut et al. 2016; Doménech et al. 2018).

Three main reasons could explain these occasional inconsistencies: first, technical issues that may affect the preprocessing of samples, PCR, and/or DNA sequencing, which could ultimately compromise sequence quality. The second reason relates to the statistical methodology used to carry out the systematic analysis. These tests may yield significantly different results depending on several factors, such as the number and length of sequences used, the number and type of species or genera studied, the alignment method, the statistical approach, or the number of applied replicates and biases caused by certain bioinformatic phenomena such as long branch attraction (Susko et al. 2021). However, the third and most important reason may lie in various biological causes, such as the accumulation of neutral mutations, their saturation, or differing nucleotide substitution rates (heterotachy) in the COI sequences of each species (Kimura 1968, 1983; Zuckerkandl & Pauling 1962; Xia 2009).

Furthermore, molecular studies of the genus have generally been conducted partially; this is by dividing analyses by geographical region (see Oeyen et al. 2014; Siriwut et al. 2015, 2016; Doménech et al. 2018). This adds to the issue, as there is no clearly defined threshold to distinguish intra-from interspecific levels for species of the genus *Scolopendra*. As a result, it is sometimes observed that the same percentage of molecular divergence between two individuals can be considered either intra- or interspecific depending on the source taken as reference (compare Oeyen et al. 2014; Siriwut et al. 2015, 2016; Doménech et al. 2018). Lastly, when comparing different species, this lack of consensus can also occasionally make it difficult to coherently reconcile morphological results with various molecular studies.

On the other hand, molecular studies on COI have generally focused on genetic positioning in trees and complemented by the percentage analysis of genetic divergences. These studies, without exception, have always been based on nucleotide sequences. However, phenotypic determination—and consequently, the external morphology—is given by the proteins translated from these and other genes, with the peptide chain carrying the “true” evolutionary weight as it determines the adaptation or elimination of individuals in a specific environment. The peptide chains of the partial COI sequence have not been studied taxonomically, yet they may offer certain advantages over nucleotide sequences. Quantitatively, amino acid chains, being two-thirds shorter than nucleotide chains, could facilitate faster reading and interpretation of different alignments. Likewise, the analysis of amino acid substitutions, where fewer changes are expected compared to nucleotide analysis (Kimura 1968, 1983), could be easier to analyze. This might translate into the use of whole numbers instead of divergence percentages obtained through more complex statistical methods subject to other statistical biases. Qualitatively, amino acid chains are more diverse than nucleotides (20 possible residues vs. only four possible bases), which allows easier discrimination of changes; making it possible to observe whether these changes are exclusive to a species or a group of species. These changes, involving electrochemical and structural modifications in the protein, can be measured, providing new differential data between sequences. Similarly, amino acid changes carry greater evolutionary and taxonomic significance than nucleotide changes, as they tend to be more inherently discriminative (Kimura 1968, 1983). Finally, protein studies would present much less bias and “noise” than nucleotide-based studies, as they would be minimally influenced by factors such as neutral mutations or substitution saturation, potentially offering molecular divergences more consistent with morphology.

Not being the main objective of this study to discern the probable causes generating the morphomolecular incongruences observed in previous articles based on mtDNA (Vahtera et al. 2012, 2013; Vahtera & Edgecombe 2014; Siriwut et al. 2016; Doménech et al. 2018), the present work analyzes for the first time the advantages and disadvantages of using translated partial COI sequences when applied to the systematics of the genus *Scolopendra*. To illustrate this, the translated partial COI fragment from a representative sample set of 21 *Scolopendra* species will be compared under this alternative molecular perspective. First, general results obtained from various bioinformatic tests will be presented, along with general conclusions for discriminating between different species of the genus. Next, as a guide, the methodology for resolving cases of difficult interpretation from an evolutionary standpoint will be outlined, exemplified by two species recently redefined by this research group — *S. spinosissima* Kraepelin, 1903 and *S. paradoxa* Doménech, 2018 — (Doménech et al. 2018; Doménech 2024). Finally, a simple summary scheme will be provided for the easy application of the aforementioned methods. The work will conclude with a discussion of the pros and cons of using these new methods and will present some taxonomic and nomenclatural observations involving some of the species studied here.

## Methods

### Pre-processing of materials, nomenclatural considerations, and image treatment

To elongate the previously trimmed terminal ends and reaffirm their suitability, the sequences of *Scolopendra spinosissima* Kraepelin, 1903 (KY888682.1), *S. paradoxa* Doménech, 2018 (MH087425.1, KY888684.1, KY888685.2, KY888686.2), and *S. subscrustalis* Kronmüller, 2009 (KY888683.2) were re-sequenced following the protocol outlined in Doménech et al. (2018).

The specific epithet of the taxon *Scolopendra hainanum* Kronmüller, 2012 ***nom. imperfect.*** is considered an original incorrect spelling (Art. 32.5 ICZN) due to grammatical gender disagreement (Art. 11.9.1.3, 31.1.2, 31.2, 32.2, and 32.5 ICZN). Therefore, the name is justifiably emended here (Art. 19.2 ICZN) and is consistently presented throughout the text as *S. hainanensis **nom. correct***.—in agreement with the species’ author, C. Kronmüller; May/June 2020, pers. comm. Likewise, the *taxon S. mojiangica* attributed to Zhang & Chi is here considered a ***nomen nudum*** (Art. 12 and 14, ICZN) as its original description is unavailable (NCBI: txid2023220). Since morphologically this species is not clearly distinguishable from other Asian congeners, this taxonomic unit is provisionally kept separate from the rest to avoid attribution bias.

The creation and editing of images (contrast and brightness adjustment, notes and references in illustrations) were carried out using Adobe Photoshop CS6.

### Bioinformatic analysis of partial COI sequences

A total of 383 COI sequences assigned to the genus *Scolopendra* were retrieved from the BOLD V4 database (Ratnasingham & Hebert 2007). The partial COI sequences were batch-translated into proteins using the getorf program from the EMBOSS tools (Rice et al. 2007), applying the invertebrate mitochondrial genetic code and discarding ORFs with fewer than 200 residues. To align the selected partial COI sequences translated into proteins, the ClustalW program implemented in BioEdit (Hall 2011) and MEGA X was used (Figure 1A, Supplementary files 1–3). The protein alignments were manually checked, discarding sequences with gaps, ambiguous residues, or those that did not align properly. Subsequently, some representative sequences from each species were selected proportionally to the number of available sequences and with the greatest possible geographical/geological distance between them. The taxonomic identity of the selected sequences was verified or modified according to current nomenclature. The divergence table of the selected protein products was generated using MEGA X (Supplementary file 4).

**Figure 1.**
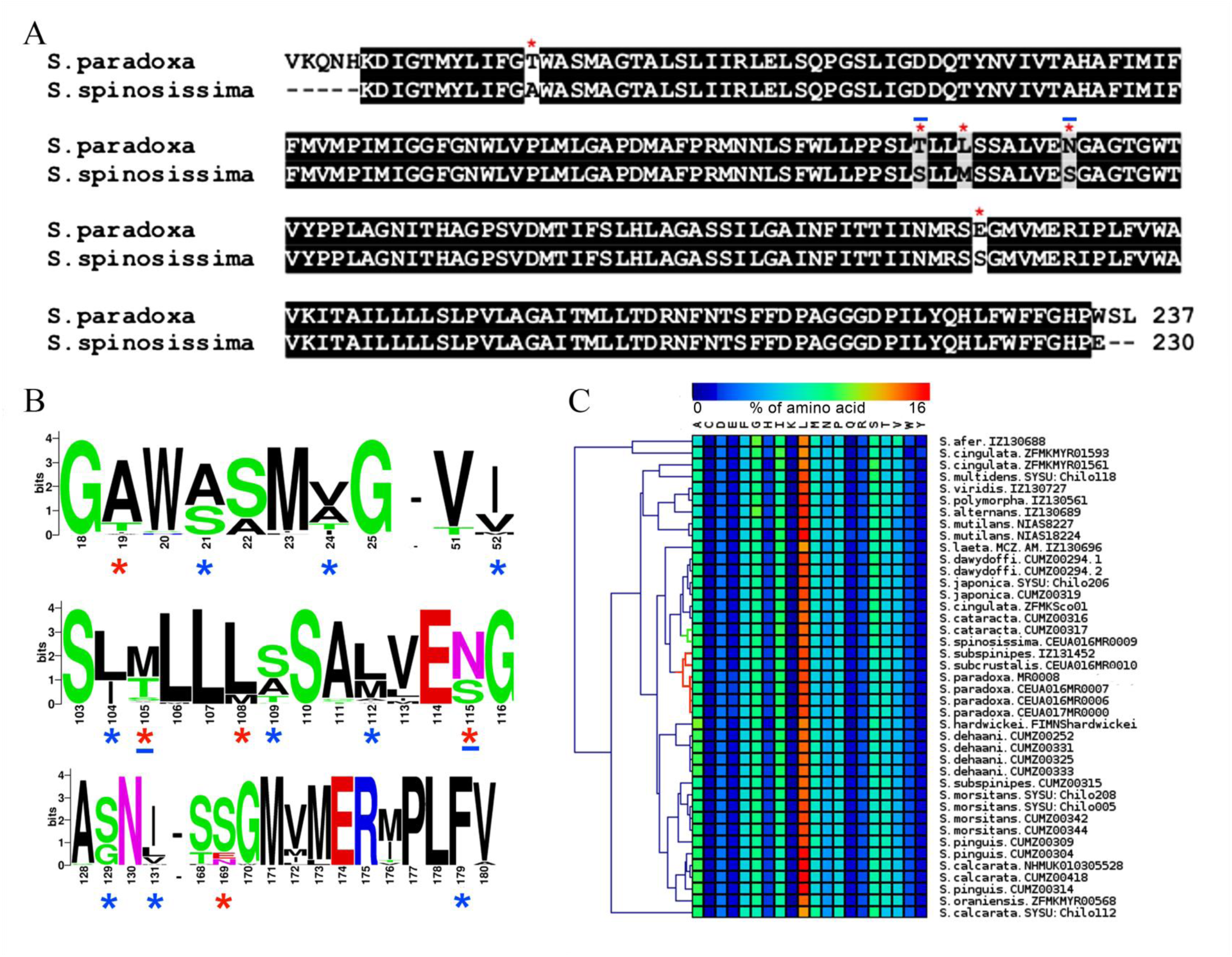
Molecular analysis of the partial COI protein sequence. **A** Pairwise alignment of the partial COI protein sequence of *S. paradoxa* / *S. spinosissima*. Identical amino acid residues are shaded in black, and residues with more than 70% similarity with the rest of the studied sequences are shaded in grey. Asterisks indicate amino acid substitutions between these two species. Blue lines mark sites where residues show high variability within the genus *Scolopendra*. **B** Logos representing 11 polymorphic residues found in the 41 partial COI protein sequences (blue asterisk). Polymorphic residues from the partial COI protein sequences of *S. paradoxa* / *S. spinosissima* are included (red asterisk). Red asterisks underlined in blue indicate polymorphic residues shared between the 22 *Scolopendra* spp. and the *S. paradoxa* / *S. spinosissima* COI protein sequences. **C** Heatmap and hierarchical clustering of amino acid frequency composition from the 45 partial COI protein sequences of *Scolopendra*. Hierarchical clustering was calculated using Pearson’s average correlation of amino acid frequency composition for each species. Colored branches highlight the close amino acid frequency relationships of *S. paradoxa* (orange) and *S. spinosissima* (green).

The amino acid substitution probability (*r*) (Table 1) was calculated using a trimmed alignment of 41 partial COI protein sequences and a substitution matrix estimated and implemented in MEGA X (Kumar et al. 2018). The sequence logos (Figure 1B) were generated using the WebLogo online tool (https://weblogo.berkeley.edu/logo.cgi). Amino acid frequencies were calculated using MEGA X, and the heat map of amino acid frequencies along with hierarchical clustering (HCL) was generated using MeV software (Multiple Experiment Viewer, www.mev.tm4.org) (Figure 1C). Amino acid sequence changes were evaluated with reference to the haplotype of the translated partial sequence belonging to the holotype of *S. paradoxa* (S. paradoxa.CEUA017-MR0000).

**Table 1.** Position and chemical proprieties of divergent amino acid residues between *S. paradoxa* and *S. spinosissima* COI protein sequence. Colum *r* is probability of substitution. The numbering of the amino acids refers to the positions indicated in the supplementary file 1. *=essential amino acids.

| Residue position | Location on highly variable regions | *S. paradoxa* |  | *S. spinosissima* |  | Probability of substitution (*r* value) |
| --- | --- | --- | --- | --- | --- | --- |
|  |  | Amino acids | Electrochemical Characteristics | Amino acids | Electrochemical Characteristics |  |
| 19 | No | T* | Polar, non-charged | A | Non-polar | 1.83 |
| 105 | Si | T* | Polar, non-charged | S | Polar, non-charged | 1.69 |
| 108 | No | L* | Non-polar | M* | Polar, non-charged | 0.47 |
| 115 | Si | N | Polar, non-charged | S | Polar, non-charged | 1.79 |
| 169 | No | E | Negatively charged;<br>Polar | S | Polar, non-charged | 0.11 |

The statistical selection of the models that best fit the amino acid substitution study was performed using the Find Best Substitution Model tool implemented in MEGA X, assuming the lowest AICc scores (corrected Akaike Information Criterion). Evolutionary analysis was carried out in MEGA X. The maximum likelihood (ML) phylogenetic tree (Guindon et al. 2010) was constructed using the mitochondrial general reversible model (mtREV) (Le et al. 2017) with 1000 bootstrap replicates (Figure 2). A consensus tree was subsequently obtained and rooted with selected outgroups. In the final stage, node support was evaluated, and genetic affinities were discussed in relation to current nomenclature and taxonomy.

**Figure 2.**
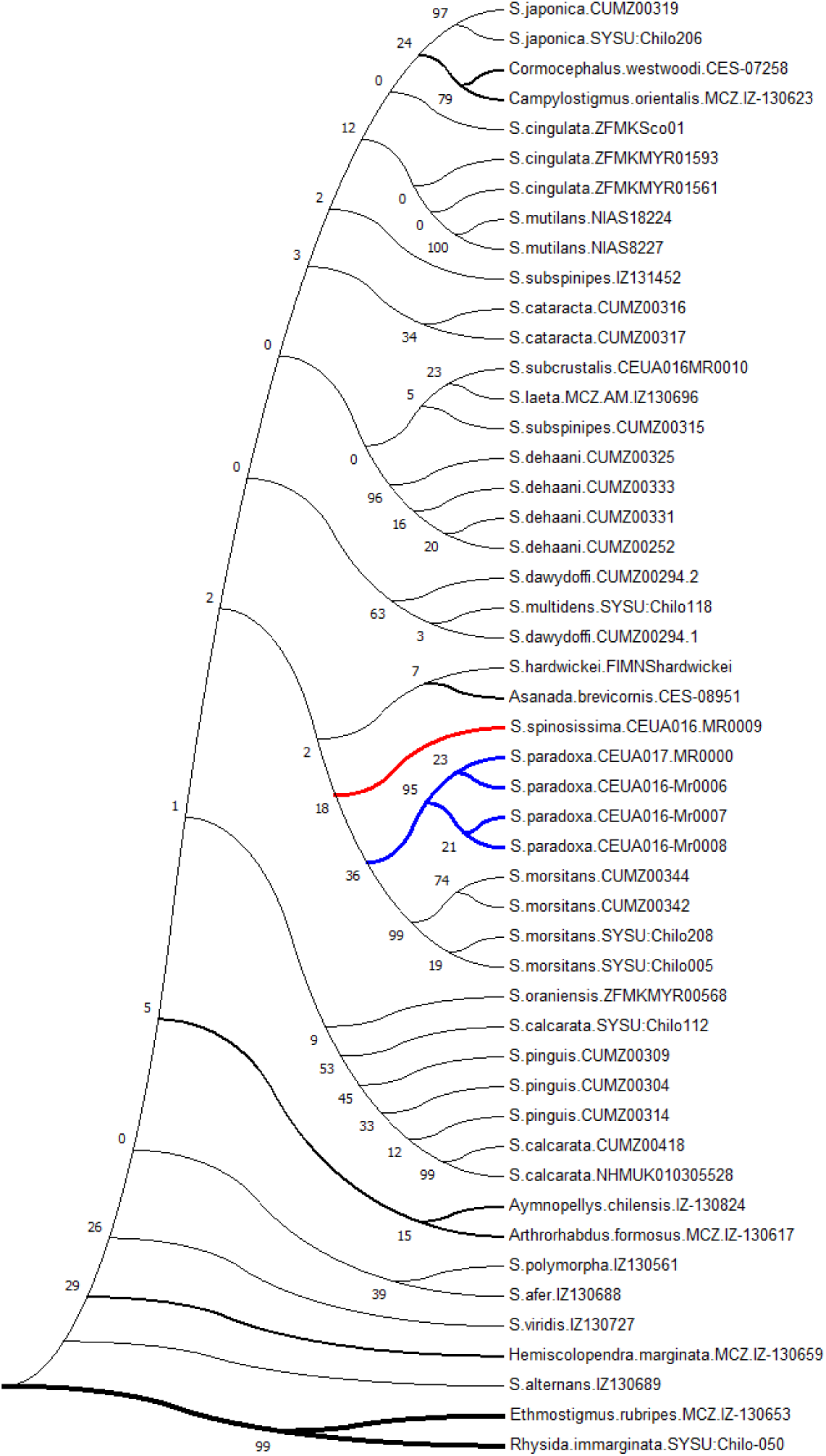
Maximum Likelihood phylogenetic reconstruction based on the protein-coding region of the partial COI gene translated into proteins, including 45 selected *Scolopendra* taxa along with six additional members of the tribe Scolopendrini and two species from the subfamily Otostigminae. Bootstrap values are shown for each node. Colored branches in the tree represent the genetic affinities of *S. paradoxa* (blue) and *S. spinosissima* (red) in relation to their congeners (thin black branches). Bold branches and thick black branches indicate the positions of other members of the *Scolopendrini* tribe and the Otostigminae species (outgroups), respectively.

The summary diagram outlining the intra- and interspecific boundaries of COI partial sequences translated into proteins for the genus *Scolopendra* was created using PowerPoint (Microsoft Office 365) (Figure 3).

**Figura 3.**
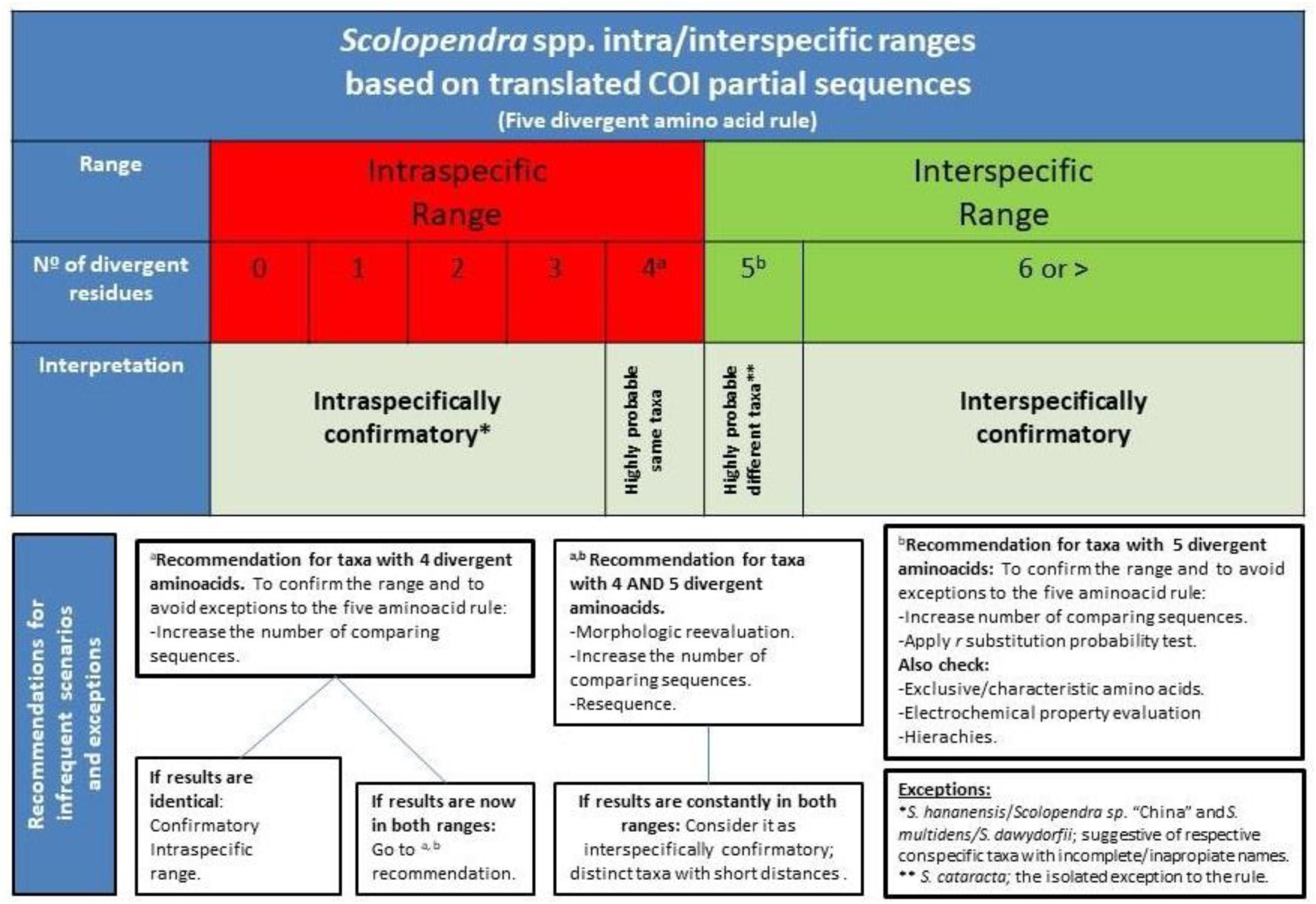
Summary scheme of the results, exceptions, and recommendations for determining intra- or interspecific differences between two or more translated protein sequences from the partial COI gene in the genus *Scolopendra*.

## Results

### Sequence Filtering

The first step in the bioinformatic analysis was to approach *bona fide* sequences and eliminate those with the evident errors. To do so, strict quality control criteria were applied to the sequences extracted from the BOLD database (see Materials section). Briefly, of the 383 sequences initially obtained, 22 sequences were discarded as they did not actually belong to the genus *Scolopendra*. Of the “29” *Scolopendra* species initially available, 14 sequences were removed for having fewer than 200 residues, including two representative species: *S*. *cf*. *amazonica* Bücherl, 1946 and *S. leki* (Waldock and Edgecombe, 2012). Additionally, another 17 sequences were eliminated for being non-translatable into a coding protein. A third filter removed 61 sequences and five more species (*S. lufengia* Kang, Liu, Zeng, Deng, Luo, Chen & Chen, 2017; *S. nigrocapitis* Zhang & Wang, 1999; *S. mojiangica*; *S. valida* Lucas, 1840; and the remaining sequences of *S. cretica* Lucas, 1853), as they yielded defective protein alignments.

Finally, from the extracted dataset, only a total of 269 sequences were deemed suitable, from which 45 sequences belonging to 21 different species were selected (Supplementary File 1).

### Partial COI protein sequences variation in the genus *Scolopendra*

Although we have found a high amino acid identity regions in our *Scolopendra* partial protein COI genes’ alignment, we have identified ten residues’ sites that present a great variability in the relative position number: 21, 24, 52, 104, 105, 112, 115, 129, 131 and 176; being also the relatives positions 249, 255, 256 and 265 other probable amino acid variable hot spots (Fig. 1B, Supplementary files 2 and 3).

Constantly, determinate amino acid combinations in the proteins’ variable fraction have being observed at intraspecific and interspecific levels. These let to establish differences between species with a quite elevate degree of confidence As a general rule, when comparing two sequences, five or more amino acid residue changes in the studied COI fragment (equivalent to 2.2% or more) indicate an interspecific range between the two amino acid chains. However, as an exception to this clear intra/interspecific division — equivalent to a mitochondrial DNA divergence gap — only 14 exceptions were observed among the 820 alignments analyzed. Therefore, this simple method of counting divergent amino acids is capable of distinguishing different species of the genus *Scolopendra* with an approximate accuracy of 98.29% (Supplementary file 4)

At the intraspecific level, all alignments uniformly showed between zero and four amino acid changes. The only exception was found in the alignment between the holotype of *S. cataracta* from Laos and a conspecific individual from China, which showed five residue substitutions. Additionally, although not counted here to avoid potential attribution/sequencing biases, two singular cases were detected far beyond these general observations: 1) *S. subspinipes* Leach, 1816, despite aligning correctly, showed either only two or nine changes respectively, depending on whether the unique CUMZ sequence was used or not; and 2) *S. pinguis* Pocock, 1891, a species under suspected cryptic speciation (Siriwut et al. 2015, 2016), which exhibited an extremely unusual range of between two and fourteen amino acid variations in the studied COI fragment (Supplementary file 4).

At the interspecific level, all alignments consistently showed five or more substitutions (hereafter referred to as the “5 divergent amino acid rule”) (Supplementary file 4). However, two different types of exceptions were detected: 1) Presumed distinct, but morphologically very similar species, with all their alignments falling within the intraspecific range (four or fewer divergent residues), such as *S. multidens* Newport, 1844 / *S. dawydoffi* Kronmüller, 2012, which consistently aligned with zero to three amino acid substitutions; and 2) Clearly distinct species whose respective alignments fell within both the intra- and interspecific range (all with between four and five substitutions): *S. cataracta* / *S. hainanensis*, *S. lufengia* / *S. dawydoffi* and *S. cingulata* when aligned with *S. lufengia* and *S. cataracta*. In the interspecific range, no divergences involving zero to three residue substitutions were observed between morphologically distinct species, leading to the conclusion that this number of changes occurs exclusively within the intraspecific range. Moreover, without exception, all morphologically distinct species with overlapping substitutions in the intra/inter range consistently had at least one alignment with four or five, or more divergent residues, depending on the sequences analyzed. This suggested that when this rare ranges overlap occurs, the analyzed taxa consistently support the presence of two distinct species (Supplementary file 4).

Moreover, upon examining the amino acid sequence alignments (Supplementary files 2 and 3), it was also found that 12 out of the 21 *Scolopendra* species had one or more unique residues in their sequence. These exclusive variations were observed consistently in species represented by multiple sequences, suggesting that the presence of such unique amino acids is, by itself, diagnostic of the species possessing them. Other amino acids were shared only by specific groups of morphologically related species, such as *S. subcrustalis* Kronmüller, 2009 and *S. laeta* Haase, 1887, or by morphologically and geographically distant species such as *S. mutilans* L. Koch, 1878, *S. polymorpha* Wood, 1861, and *S. viridis* Say, 1821, indicating a character shared by a group of species. When these group-characteristic but not species-exclusive amino acids were found, they were also observed to be useful for distinguishing species—especially when the taxa were not morphologically and/nor geographically related.

On the other hand, in terms of amino acid composition, the different translated partial COI protein sequences also showed significant similarity among *Scolopendra* species (Figure 1C). However, slight variations in composition were observed, which made it possible to establish hierarchical groupings and, in turn, distinguish species by positioning them in a way that differs from the phylogenies (Figure 2). By this method, only some species showed non-monophyletic groupings.

At the phylogenetic level, the tree inferred based on protein chains (Figure 2) showed similar results in species placement when compared to mtDNA trees (check Siriwut et al. 2015, 2016; Oeyen et al. 2014; Doménech et al. 2018). However, branch positioning was more sensitive to amino acid variations, while node supports were correspondingly weaker in protein-based phylogenetic reconstructions. Ultimately, it was found that trees based on COI protein-coding sequences were somewhat less useful for establishing phylogenies than those based on mtDNA (see Siriwut et al. 2015, 2016; Oeyen et al. 2014; Doménech et al. 2018).

At the taxonomic level, the molecular results showed a significant number of amino acid substitutions (>4) between *S. mutilans* and its closest relative, *S. subspinipes*, reaffirming that these entities are not conspecific taxa, contrary to what has been suggested in recent literature (Vahtera et al. 2013; Kang et al. 2017; Han et al. 2018) (Supplementary File 4). Conversely, the findings by Kang et al. (2017) which reported no molecular differences between *S. subspinipes* and *S. hainanensis*, are not supported here (residue differentiation >4; Supplementary File 4), as our results were consistent with them being distinct species, aligning with their clear morphological differences (Kronmüller 2012; Siriwut et al. 2016).

In parallel, experimental alignments (unpublished), which include relatively short but partially well-conserved sequences, revealed that the morphological similarity between *S. morsitans* Linnaeus, 1758 and *S. cf. amazonica* is strongly supported at the molecular level, suggesting a likely synonymy (residue substitution = 1). Meanwhile, species that appear morphologically similar, such as *S. japonica* and *S. negrocapitis* Zhang & Wang, 1999, as well as *S. mojiangica*, produced inconclusive results (three to five amino acid substitutions), showing the unsolved taxonomical and nomenclature situation for these cases.

### COI protein sequence variations between *S. paradoxa* and *S. spinosissima*, and final considerations

The analysis of COI protein sequence variations was conducted using the DNA sequences of *S. paradoxa* and *S. spinosissima* obtained in Doménech et al. (2018) (Supplementary File 1). We found that the protein sequences derived from the holotype and paratypes of *S. paradoxa* encoded exactly the same protein sequence (Supplementary Files 2 and 3). Therefore, only one consensus sequence of *S. paradoxa* was used for the following analyses. Figure 1A shows the pairwise alignment of the partial COI protein sequences of *S. paradoxa* and *S. spinosissima*. As shown in the alignment, the partial COI proteins displayed a high degree of identity (97.8% of aligned amino acids in Supplementary Files 2 and 3). However, we identified five amino acid residue differences between the two species: 19T>A, 105T>S, 108L>M, 115N>S, and 169E>S (Figures 1A–B, Table 1). Of these five substitutions, two—105T>S and 115N>S—were located in regions of high variability observed across the sequences (Supplementary File 2). The variations 19T>A, 108L>M, and 169E>S involved relevant changes in the physicochemical properties of the amino acids (Table 1), which could suggest changes in the structure, function, and adaptation of the resulting COI protein.

To ensure that the evolutionary process between these two species—of which few sequences are available—is not random or an exception to the “five amino acid rule,” we calculated the substitution probability of these five residues using a total of 41 partial COI protein sequences of *Scolopendra* (Table 1). We found that substitutions 19T>A, 105T>S, and 115N>S had a substitution probability greater than one, suggesting strong positive selection (adaptive or diversifying) at these three amino acid positions (Table 1). In contrast, the substitutions at residues 108L>M and 169E>S showed substitution probabilities below one, suggesting moderate and strong negative (purifying) selection, respectively (Table 1). These data support the conclusion that the observed amino acid divergence cannot be explained by genetic drift alone and highlight the selective pressures involved in the diversification of these two species.

Among all the *Scolopendra* species represented in this study, *S. paradoxa* consistently and exclusively exhibited a glutamic acid (E) at relative position 169, making this amino acid a standalone diagnostic character for the species (Supplementary File 3). As a group, *S. paradoxa*-*S. spinosissima* did not exhibit unique group-specific amino acids that differentiated them from the rest of the species. However, *S. spinosissima* shared characteristic amino acids with *S. calcarata* Porat, 1876, while *S. paradoxa* shared group-specific amino acids with *S. laeta*—none of which were found together in any other species.

In the hierarchical classification of amino acid frequencies (Figure 1C), *S. paradoxa* and *S. spinosissima* were grouped into non-sister sections, suggesting the separation of these species by the use of this method. Similarly to this, the inferred protein phylogenetic reconstruction showed *S. paradoxa* and *S. spinosissima* as close but not strictly sister species. *Scolopendra paradoxa* displayed strong monophyletic node support including its four sequences, whereas the position of *S. spinosissima* remained weakly supported—precisely as reported in previous literature (Doménech et al. 2018).

In line with earlier findings (Doménech et al. 2012), all these showed molecular analyses concerning *S. spinosissima* and *S. paradoxa* once again conclude that these taxonomic entities are objectively and definitively two distinct species, separated by evolutionary selective pressures rather than by chance.

Finally, to facilitate species discrimination within *Scolopendra* using this alternative molecular method based on partial proteins translated from COI fragments, we provide a summary diagram of the results obtained, including noted exceptions and recommendations for particular situations (Figure 3).

## Discussion

In this study, the translated partial COI gene has been confirmed as a valuable biomarker for the identification and differentiation of species within the genus *Scolopendra*. The analysis of amino acids in the partial COI protein fragment showed that sequences belonging to different species differ by five or more residues. However, some exceptions to this “rule” were detected. To clarify most of these ambiguous inter/intraspecific cases, additional analysis of exclusive and characteristic divergent amino acids—their frequency, electrochemical properties, and especially the *r* substitution probability test—proved to be useful tools for resolving these exceptional scenarios.

Compared to traditional divergence studies based on mtDNA percentage differences, the intra/interspecific discrimination of amino acid substitutions expressed as natural numbers proved to be more easily interpretable, computationally less demanding, and less affected by methodological and biological variation. Likewise, the *r* substitution probability test showed a high degree of confidence in resolving complex cases of taxon differentiation, providing measurable insights into the evolutionary history of these species. On the other hand, the phylogenetic tree inferred from amino acid chains, while yielding good results on its own, also proved to be no more accurate than phylogenetic reconstructions based on mtDNA (Figure 2). In this case, the shorter and more similar sequences, along with the statistical weight of any amino acid substitution, made this type of analysis—though robust—more prone to fluctuations than the more “tolerant” trees built from mtDNA data. Meanwhile, heat map and hierarchical clustering, although useful in distinguishing species by amino acid composition in most cases, were not as precise as divergence studies, but still served as valuable supporting tools for the overall analysis. In contrast, the study of exclusive or group-characteristic amino acids proved to be significant for the separation and discrimination of *Scolopendra* species. However, this latter method should be interpreted with caution in species with a low number of sequences analyzed and/or a restricted geographic distribution.

Proportionally, it has been shown that interspecific mtDNA divergence percentages do not correspond to those of amino acids (i.e., compare the divergences of *S. spinosissima* and *S. paradoxa*, with respective differences of 19% for mtDNA and 2.2% for the coding protein; see Supplementary File 4 and Doménech et al. 2018). This result aligns with Kimura’s neutral theory (1983), which states that most DNA mutations do not affect codon translation. Therefore, compared to most nucleotide changes, an amino acid substitution is much more influential on the structure and function of the resulting protein, and consequently, on species decline or differentiation. Thus, due to its inherently lower susceptibility to the random effects of evolution, a species differentiation method based on proteins, such as the one proposed here, could represent an alternative to mtDNA-based methods, offering certain interpretative advantages and a high level of reliability.

On the other hand, from a general perspective, it is noteworthy that approximately one-third of the available COI sequences of Scolopendra exhibited various issues that hindered proper alignment. The intrinsic difficulty of COI sequencing—which includes artifacts such as nuclear mitochondrial DNA segments (NUMTs) (Rogers & Griffiths-Jones 2012)—combined with human-related factors like species misidentification, statistical errors, and biological elements such as the accumulation of neutral mutations, excessive substitution saturation, or the phenomenon of heterotachy (Kimura 1968, 1983; Zuckerkandl & Pauling 1962; Xia 2009), may ultimately disrupt molecular analysis. All of these elements should be evaluated to improve the quality of future analyses aimed at species differentiation. Furthermore, aside from the sequences reviewed directly in this study, the quality of the remaining protein sequences filtered using different methods can only be approximated here, and cannot be guaranteed with certainty. For this reason, some of the “irregular” cases tentatively treated as exceptions in this study (see Results section) may, in fact, be explained by the aforementioned errors. A higher standard of rigor and standardized filtering applied prior to sequence publication could be key to improving the overall quality of COI databases.

In the species-level analysis, our study found that some sequences of *S. cataracta* exhibited the highest molecular variability within the genus, consistent with the findings of Siriwut et al. (2015), making it the only intraspecific exception to the “five amino acid rule.” However, despite passing the selection filters, this exception could potentially be linked to an error, given the geographic-molecular discrepancies observed among the available sequences of this species, thus requiring further study. On the other hand, the morphologically very similar S. multidens and S. dawydoffi (Siriwut et al. 2016) also could not be separated using this method, which might suggest that the minor morphological differences actually represent intrinsic variability within a single taxon. The possible synonymy of these two species should be confirmed through a study of the type specimens. Likewise, the sequences of *S. subspinipes* and *S. pinguis* should be re-examined due to the excessively high intraspecific distances observed in this study. Additionally, the validity of the few sequences available for *S. cingulata*, *S. cataracta*, and *S. lufengia* should also be assessed, as their placement falls ambiguously within either the intra- or interspecific range. If no errors are confirmed in these few cases, such deviations should be interpreted solely as a reflection of the progressive evolutionary effect in certain specimens—representing a transitional and certainly exceptional stage relative to the divergence ranges clearly defined by the five amino acid rule.

Regarding *S. spinosissima* and *S. paradoxa*, the comparison of the translated COI ortholog gene revealed five clear and consistent differences in their amino acid composition and in certain electrochemical properties (Figure 1A and B, Table 1, Supplementary Files 2 and 3). Moreover, the *r* substitution probability analysis rejected the null hypothesis (genetic drift) for all amino acid substitutions and confirmed that none of them could be explained by any random process. All these genetic results provide evidence of a speciation process driven by selection between *S. spinosissima* and *S. paradoxa*, findings that further support the morphological differences and taxonomic separation of both species (Doménech et al. 2018, Doménech 2024).

The genetic results presented here represent a step forward in the distinction of *Scolopendra* species. Although consistent, the conclusions of this analysis were based on representative sequences from approximately 20–25% of the described species within the genus, and are therefore subject to further refinement. Revisiting the questionable and excluded sequences, along with incorporating more sequences from both these and other unexamined taxa, could enhance the accuracy of this new and simple methodology aimed at improving the molecular differentiation of *Scolopendra* species.

## Abbreviations

**Institutions**
BMB-DENR: Biodiversity Management Bureau - Department of Environment and Natural Resources, Philippines.
CEUA: Colección Entomológica de la Universidad de Alicante San Vicent del Raspeig, Alacant, Spain.
PAE: Philippines Association of Entomologists, Inc., Philippines.
PNM: Philippine National Museum, Philippines.
PNU: Philippine Normal University, Philippines.
WRD-DENR: Wildlife Resources Division - Department of Environment and Natural Resources, Philippines.

**Molecular Section**
A: alanine;
COI: cytochrome *c* oxidase;
E: glutamic acid;
L: leucine;
M: methionine;
N: asparagine;
S: serine;
T: threonine.

**Others**
ICZN: International Code of Zoological Nomenclature.

## Acknowledgments

The authors sincerely thank S. Rojo (UA) for his excellent and constructive advice during the preparation of this manuscript. Additional thanks go to J. B. Warren and E. C. Jarvis for their work correcting the spelling and grammar of the text. Recognition is also extended to S. A. Yap, G. D. Sapin, G. A. Lantican, J. J. Apolinario (PAE), P. A. C. Buenavente (PNM), and V. San Juan (PNU), and especially to M. C. Amaro Jr. (BMB-DENR) and P. Bumanglag (WRD-DENR) for their invaluable bureaucratic support and for authorizing the use of *Scolopendra* sequences from the Philippines (Agreement of October 18^th^, 2023 – https://bmb.gov.ph, see also GenBank). In accordance with the Philippine authorities, the undersigned authors of this article wish to explicitly emphasize that access to any type of biological resources in this nation always requires appropriate research permits from the DENR.

## Supplementary files

**Supplementary file 1.**
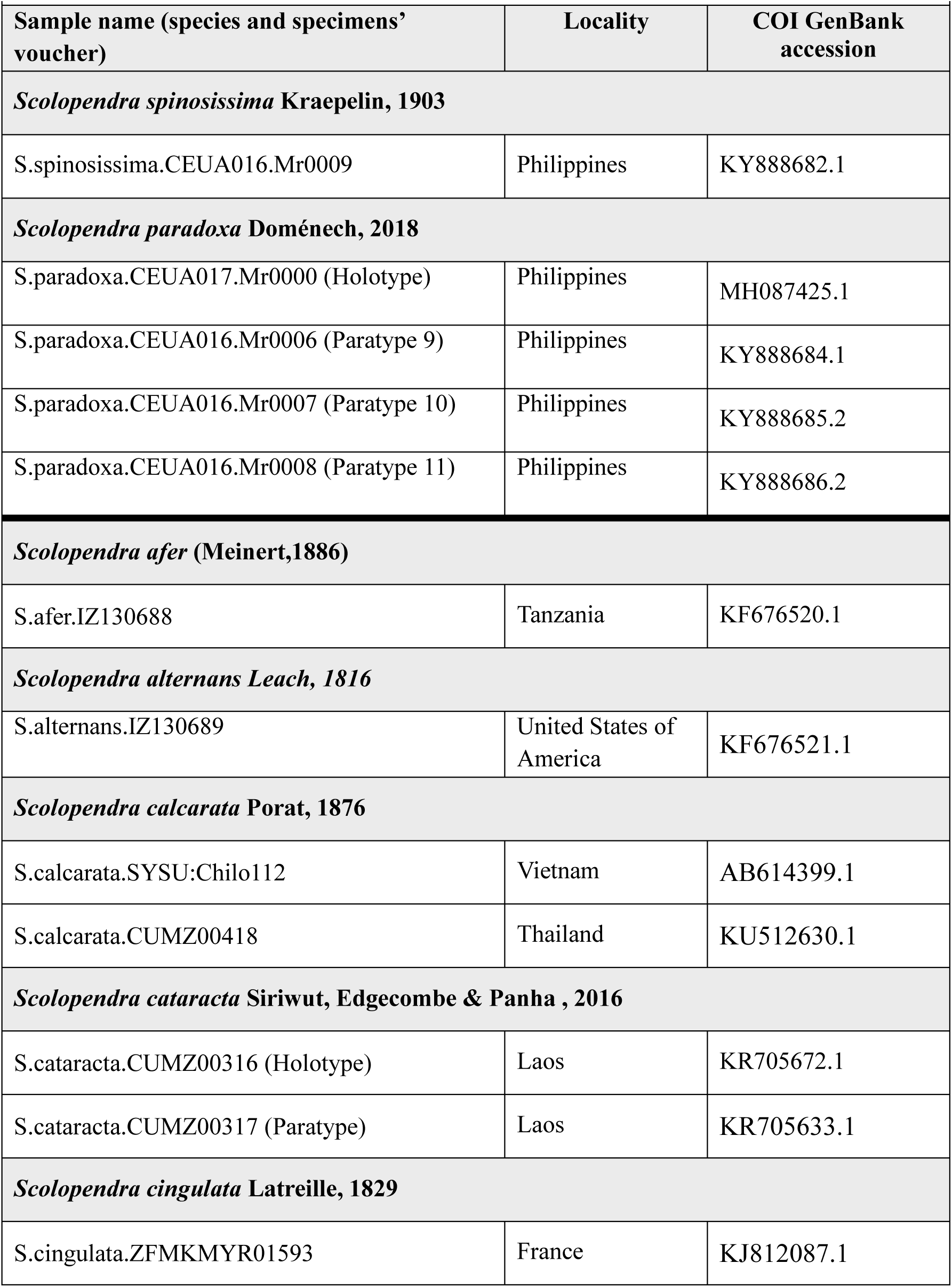
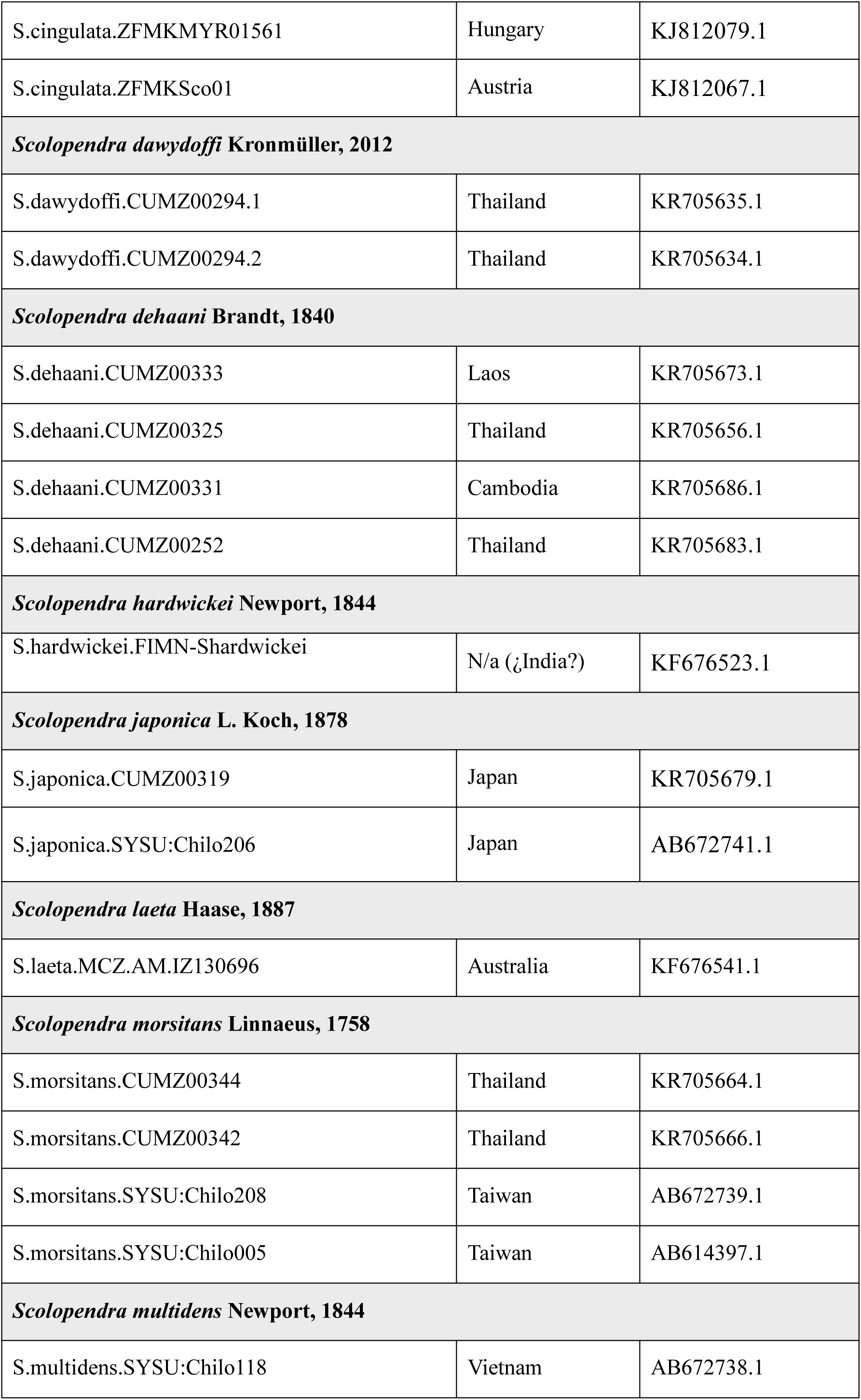
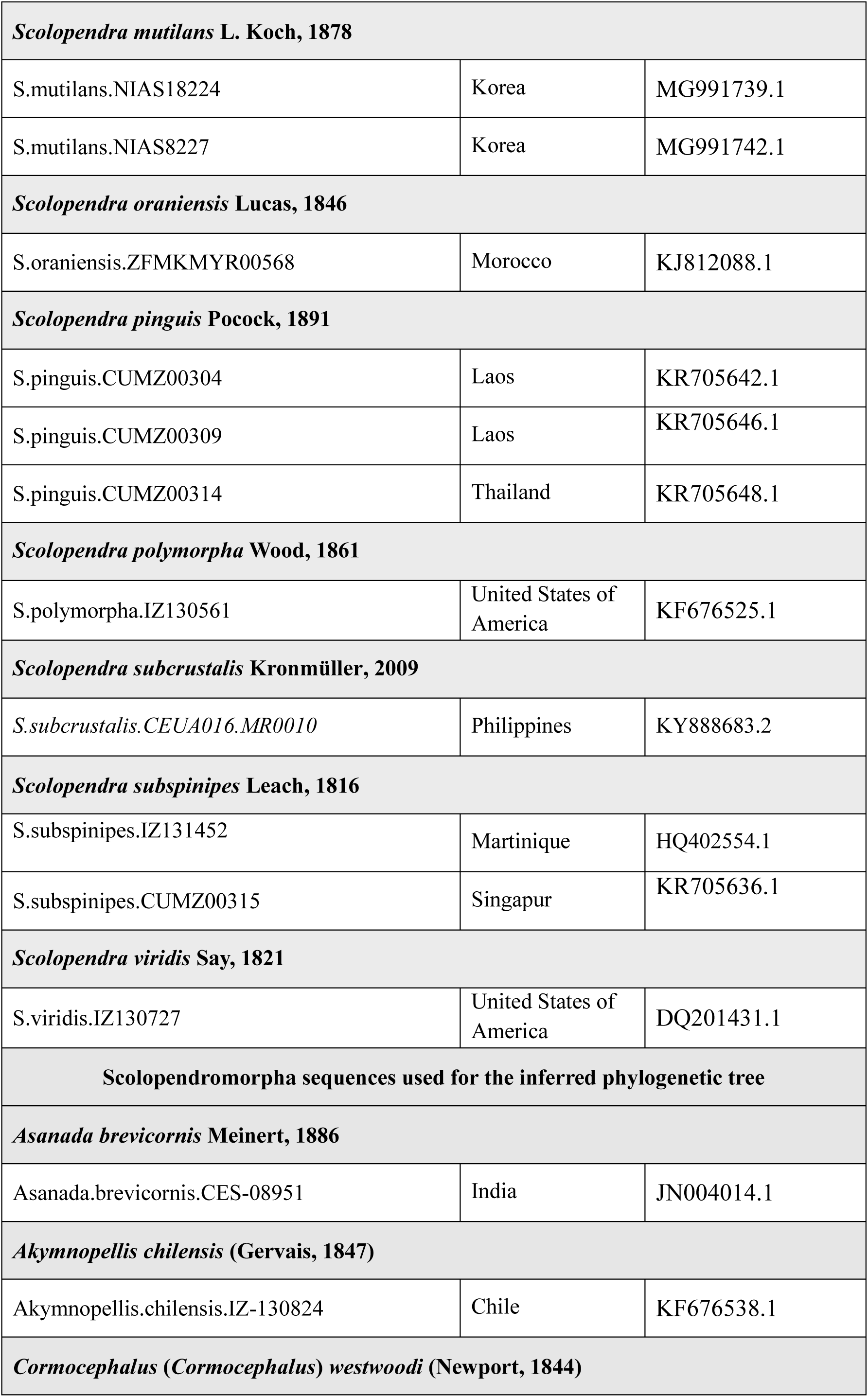
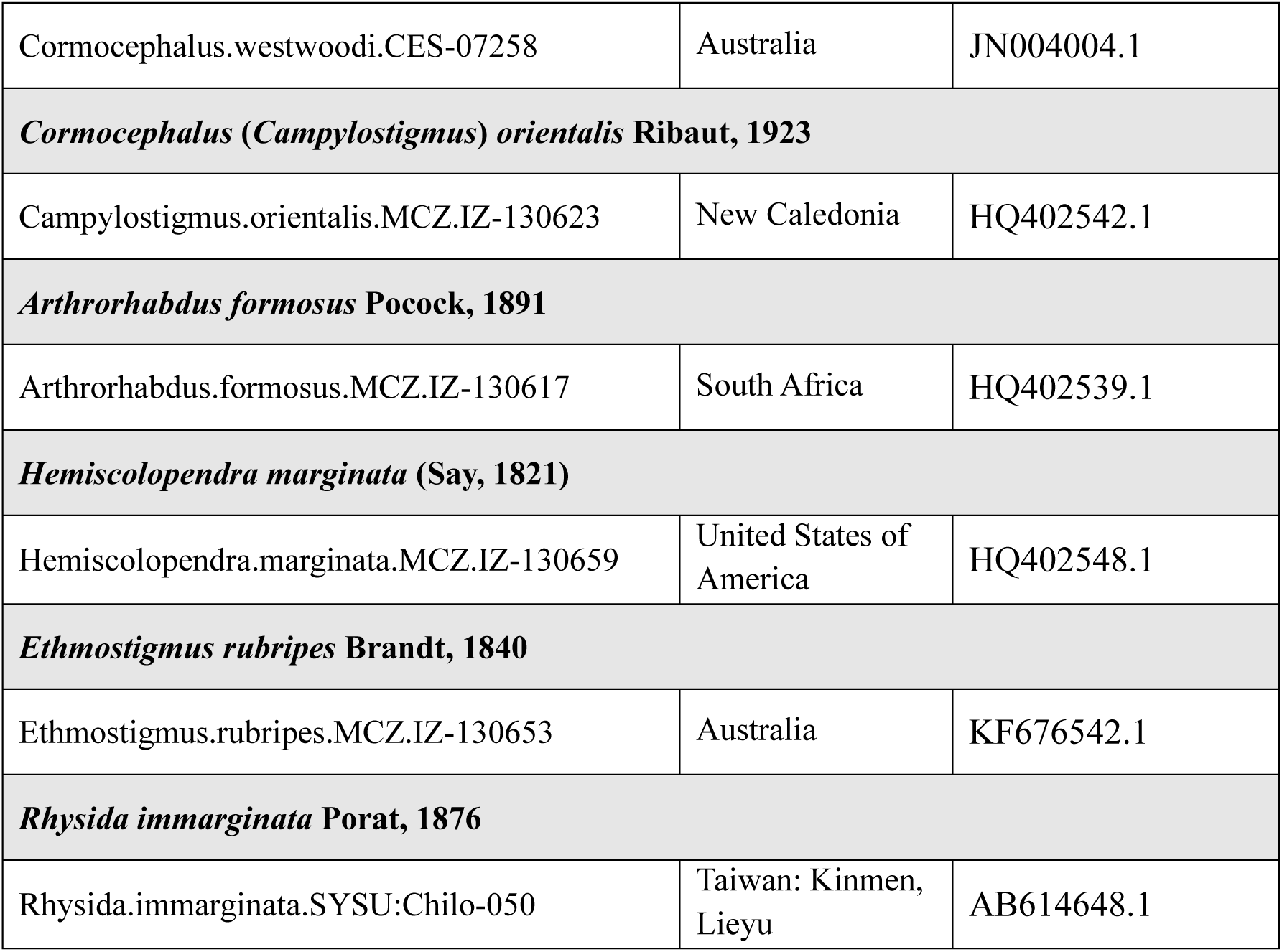
Summary of the 45 COI sequences from *Scolopendra* species, along with eight from other Scolopendromorpha, obtained from the Barcode of Life Data System (BOLD) and used in the molecular analyses. Each sample includes the institutional abbreviation of the voucher, type status, voucher code, locality, and GenBank accession number for the selected partial COI gene sequence.

**Supplementary file 2.**
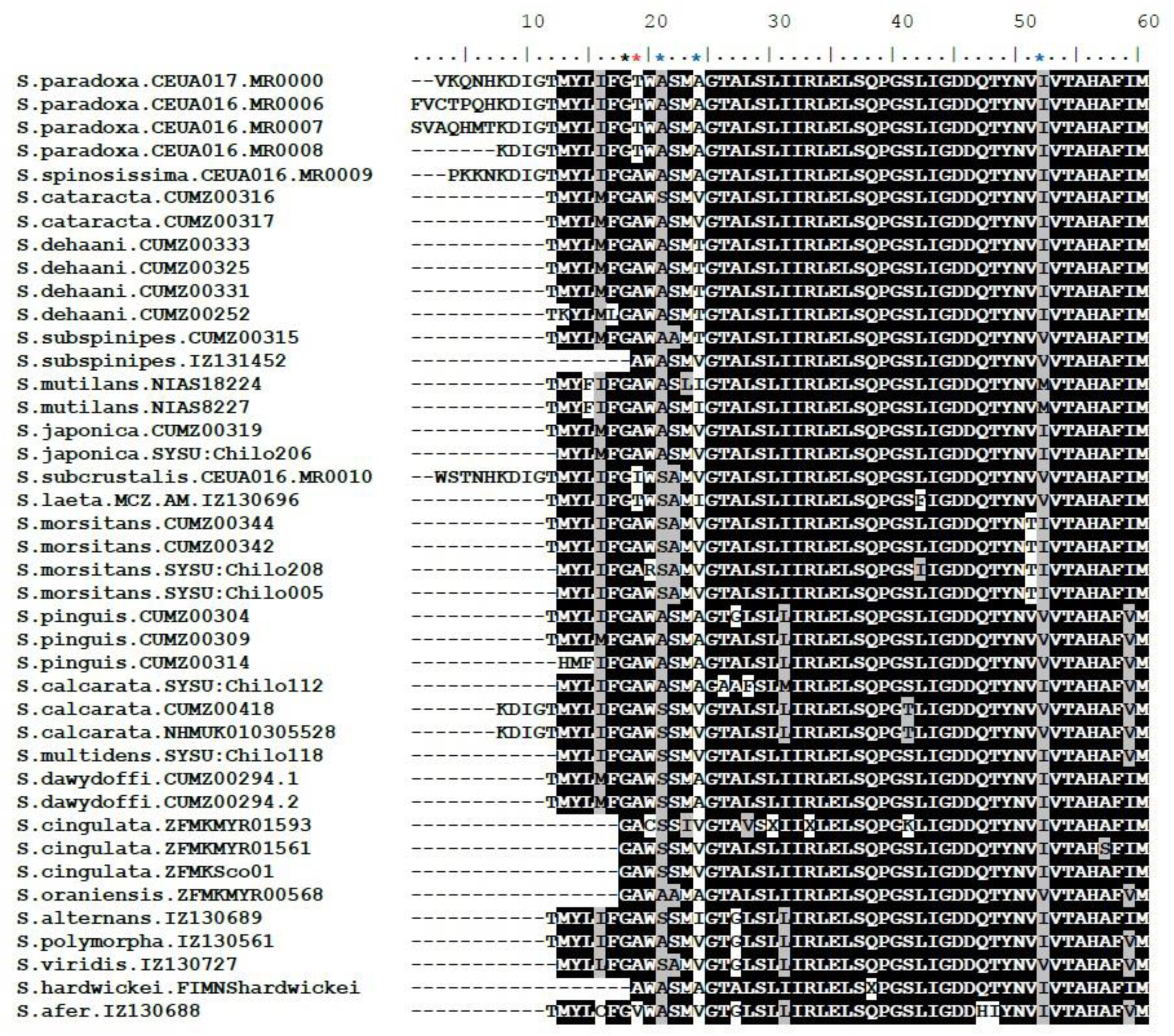
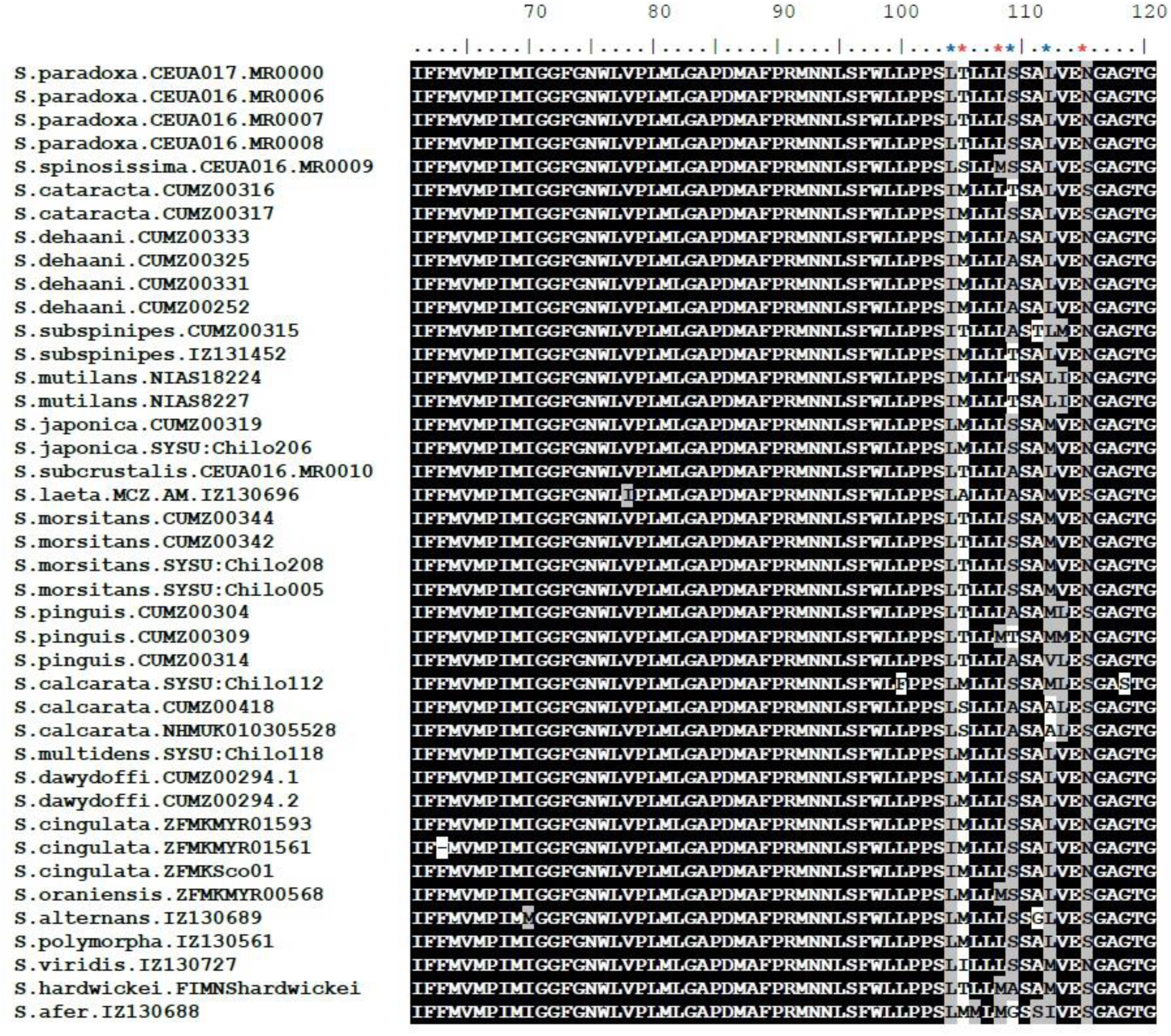
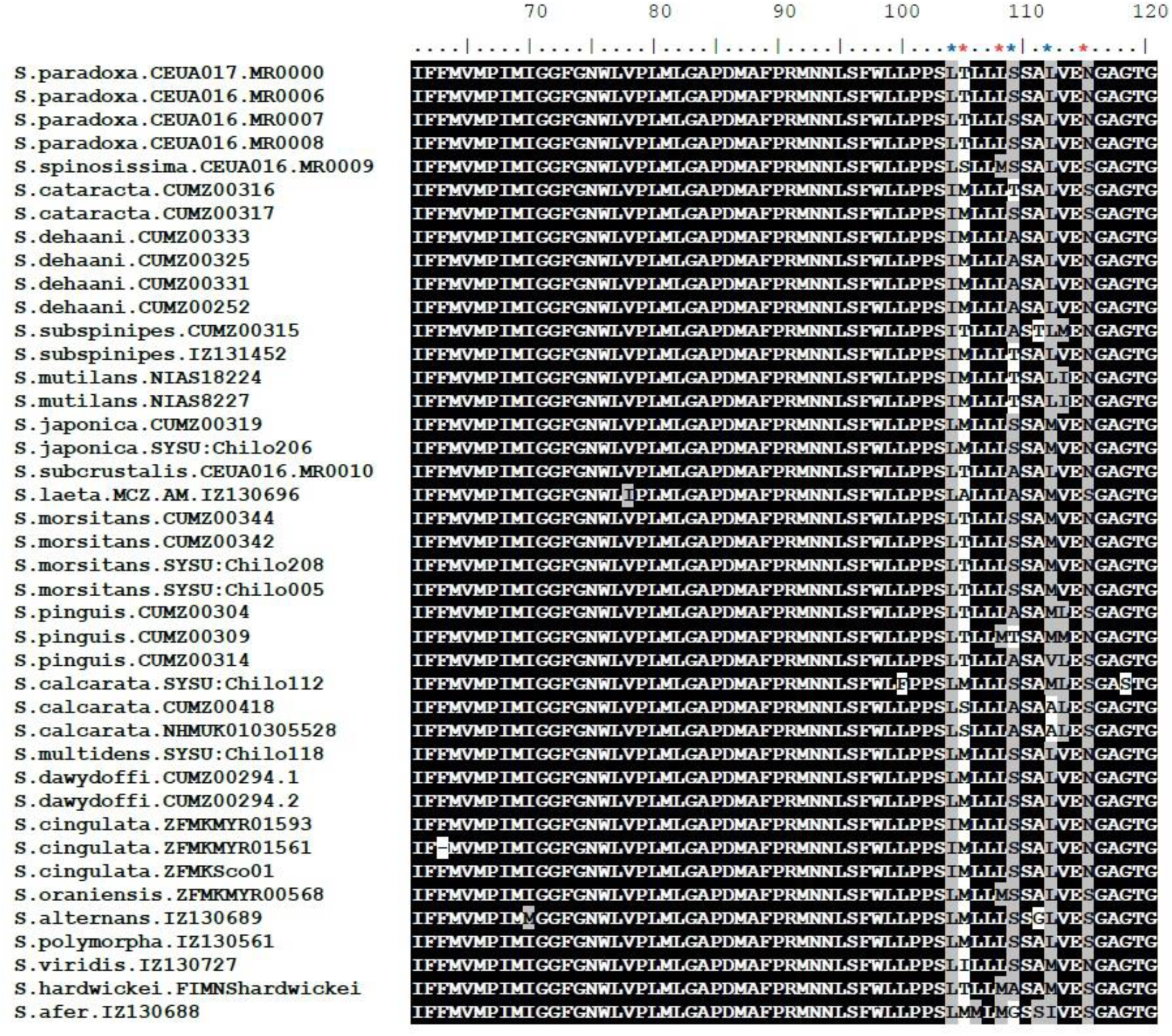
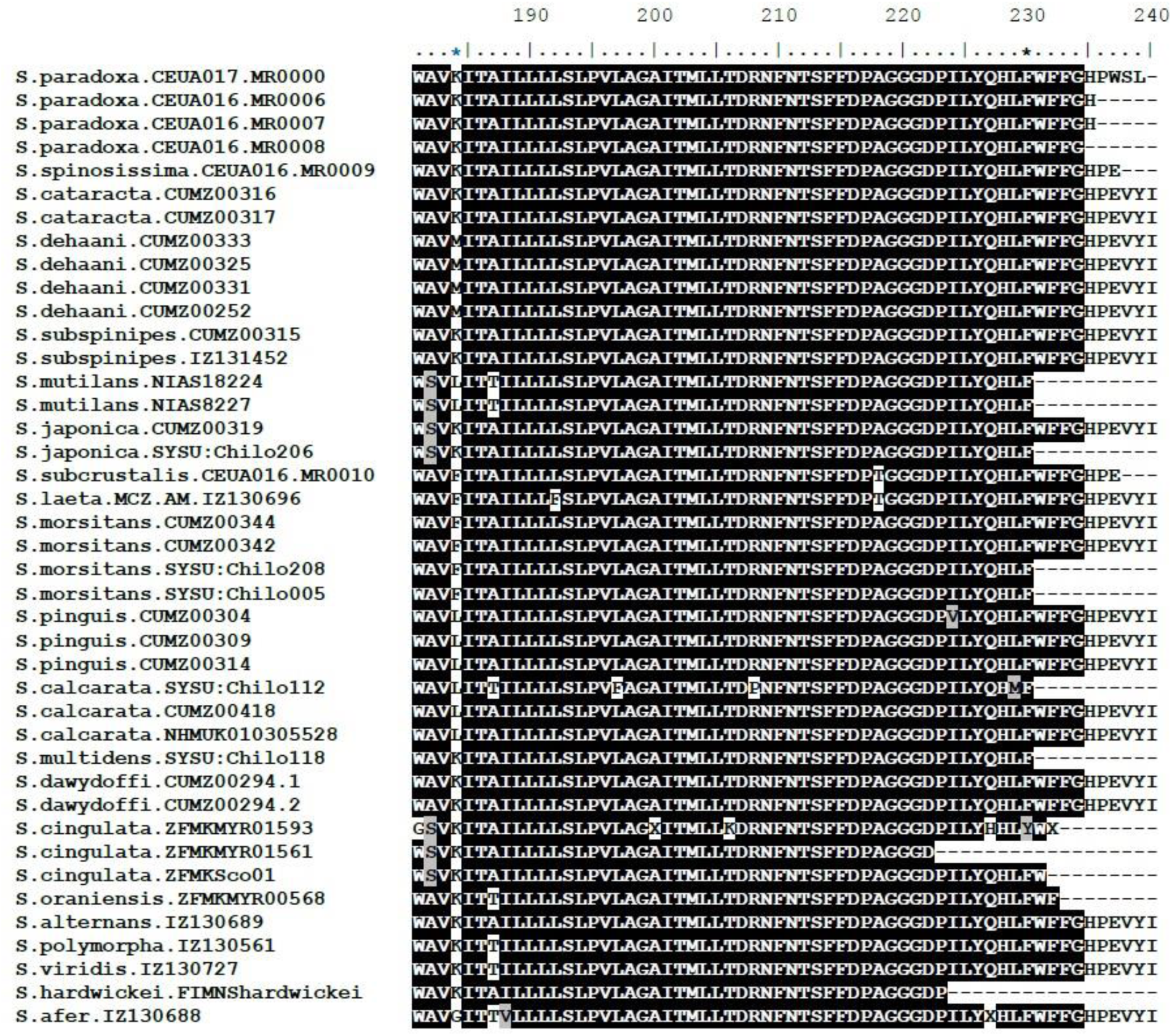
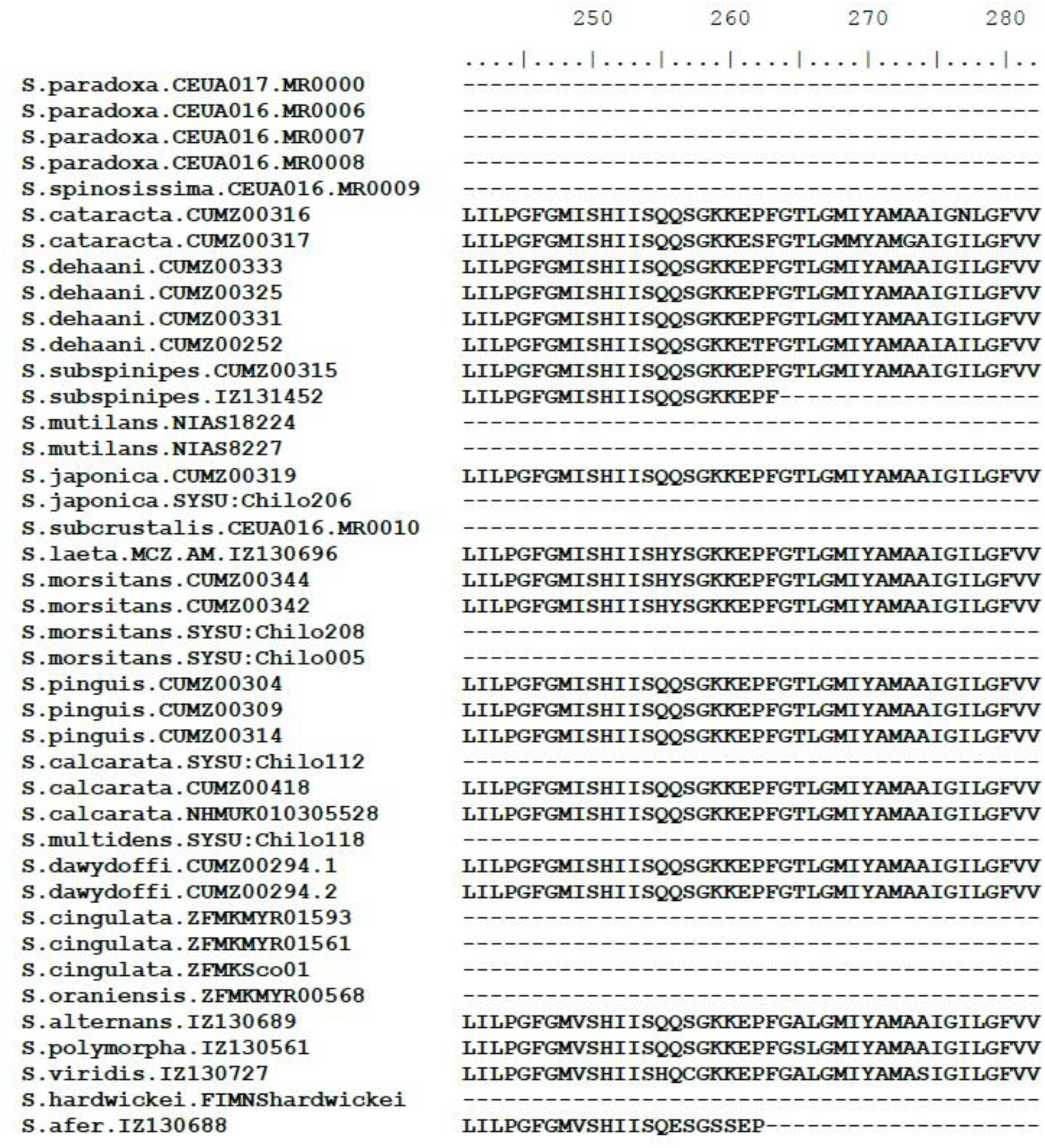
Alignment of the translated amino acid sequence from 45 partial DNA sequences of the COI gene. The alignment was performed using the ClustalW algorithm in the BioEdit package. Black asterisks indicate the first and last amino acid positions of the trimmed alignment used in the molecular analysis. Red asterisks indicate the five differing amino acid residues between the COI protein sequences of *S. paradoxa* and S. spinosissima. Blue asterisks indicate variable amino acid residues in the partial COI sequence within the genus *Scolopendra*. Black boxes mark conserved amino acid sites. White boxes indicate variable sites with <70% similarity, while gray boxes indicate sites with ≥70% similarity.

**Supplementary file 3.**
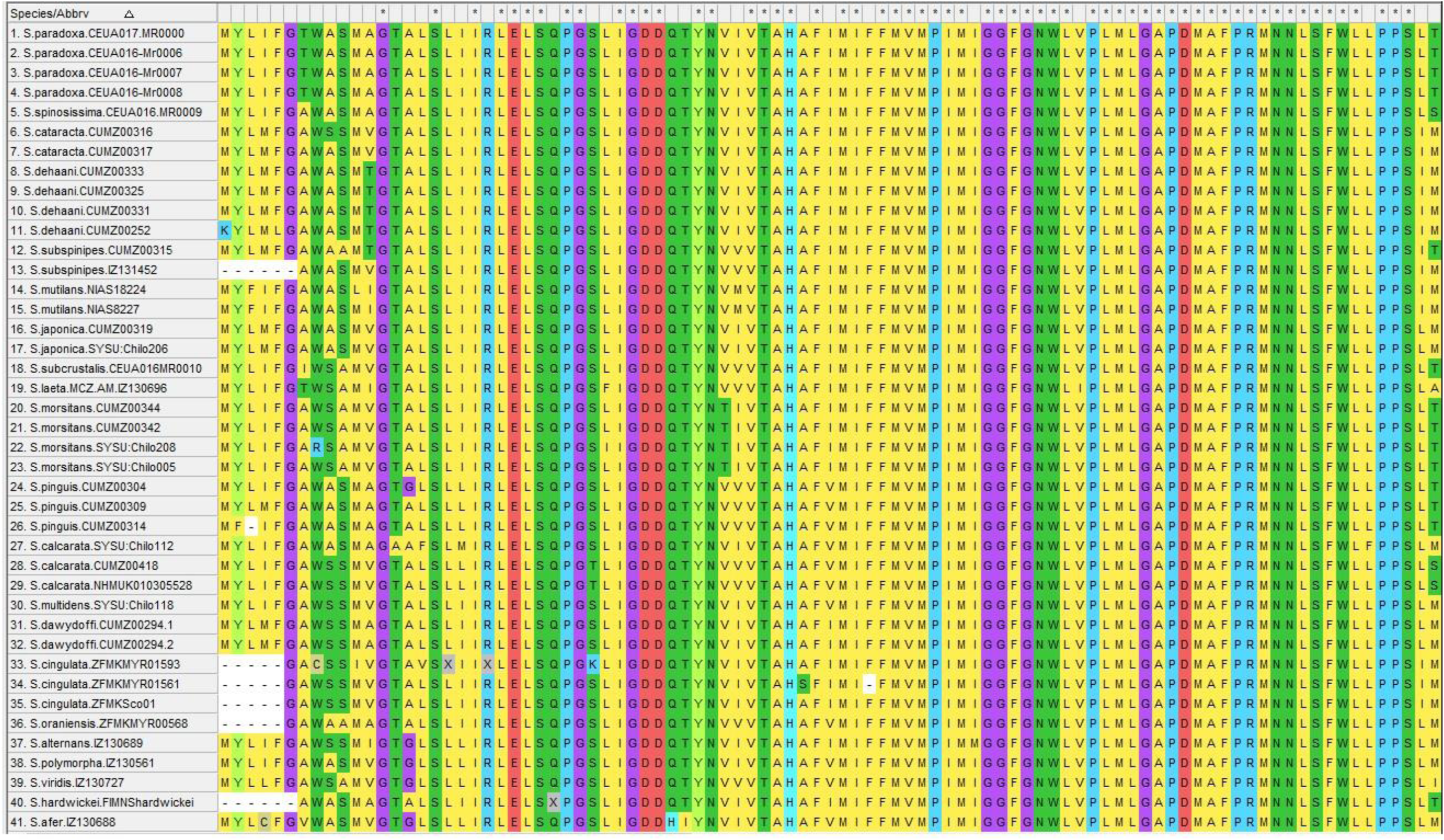
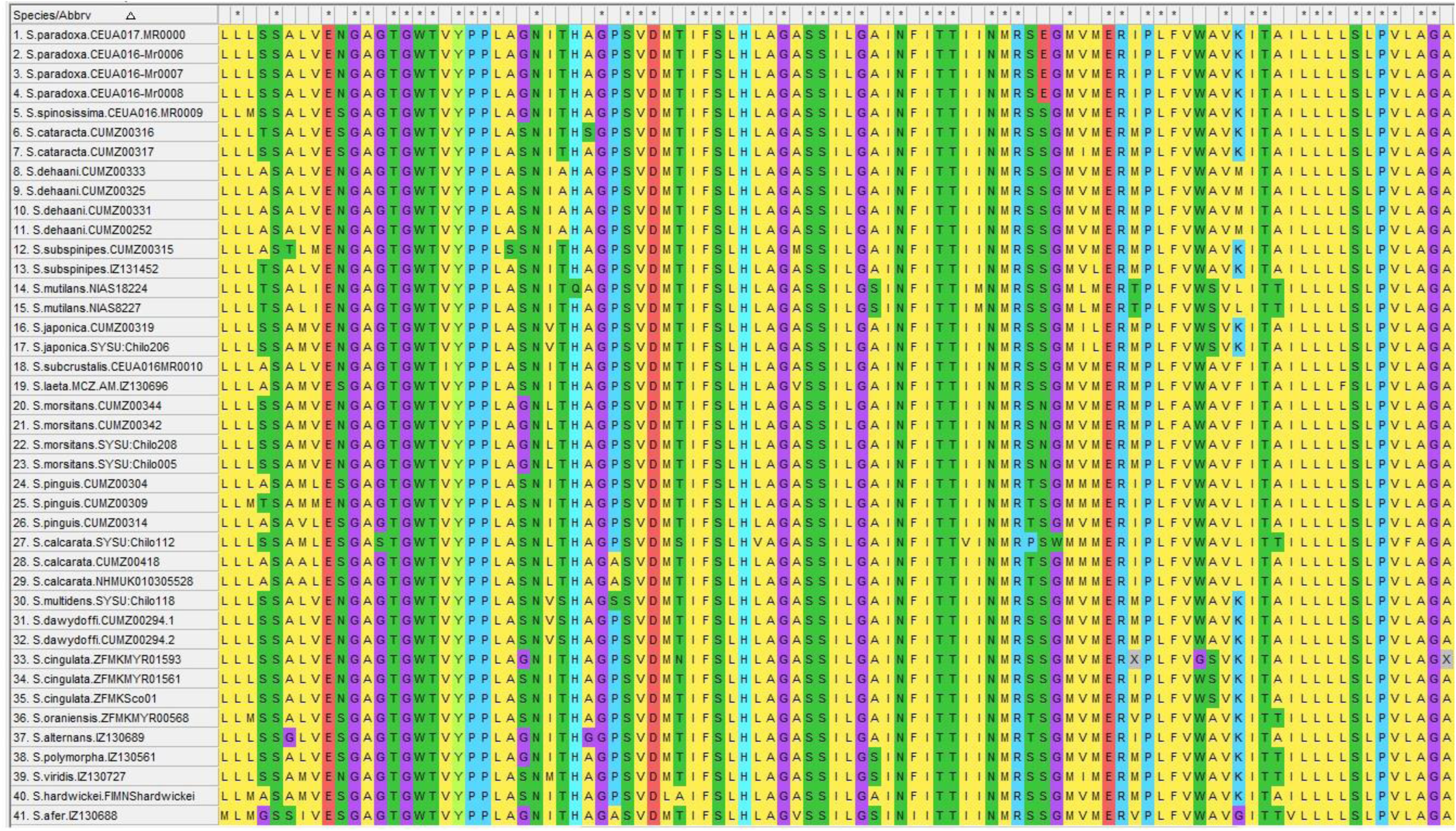
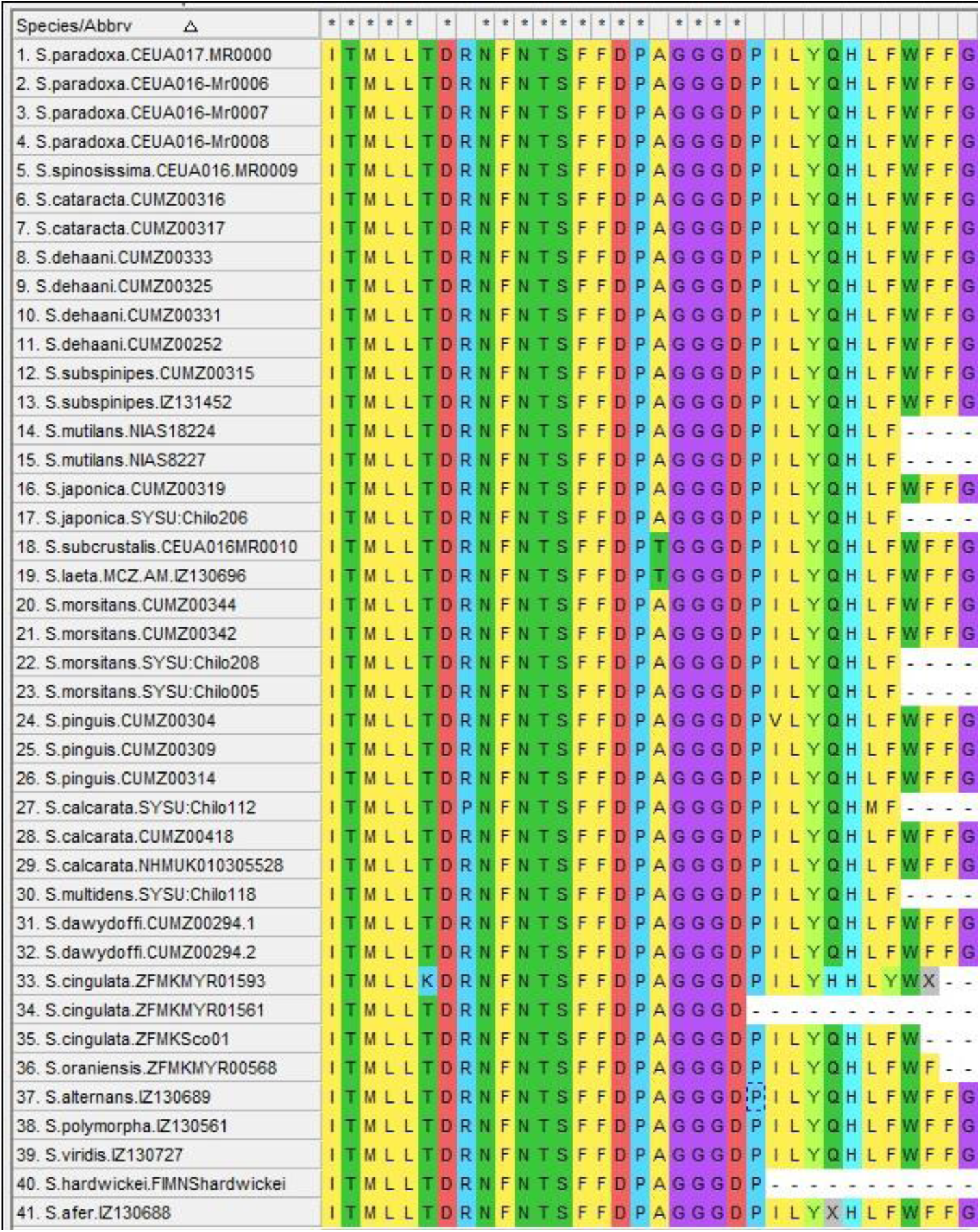
Alignment of 45 amino acid sequences from the partial COI gene belonging to 22 different species of the genus *Scolopendra*. Asterisks indicate residues common to all studied species. The color coding helps identify amino acids that are unique to a species and characteristic of a group of species.

**Supplementary file 4.**
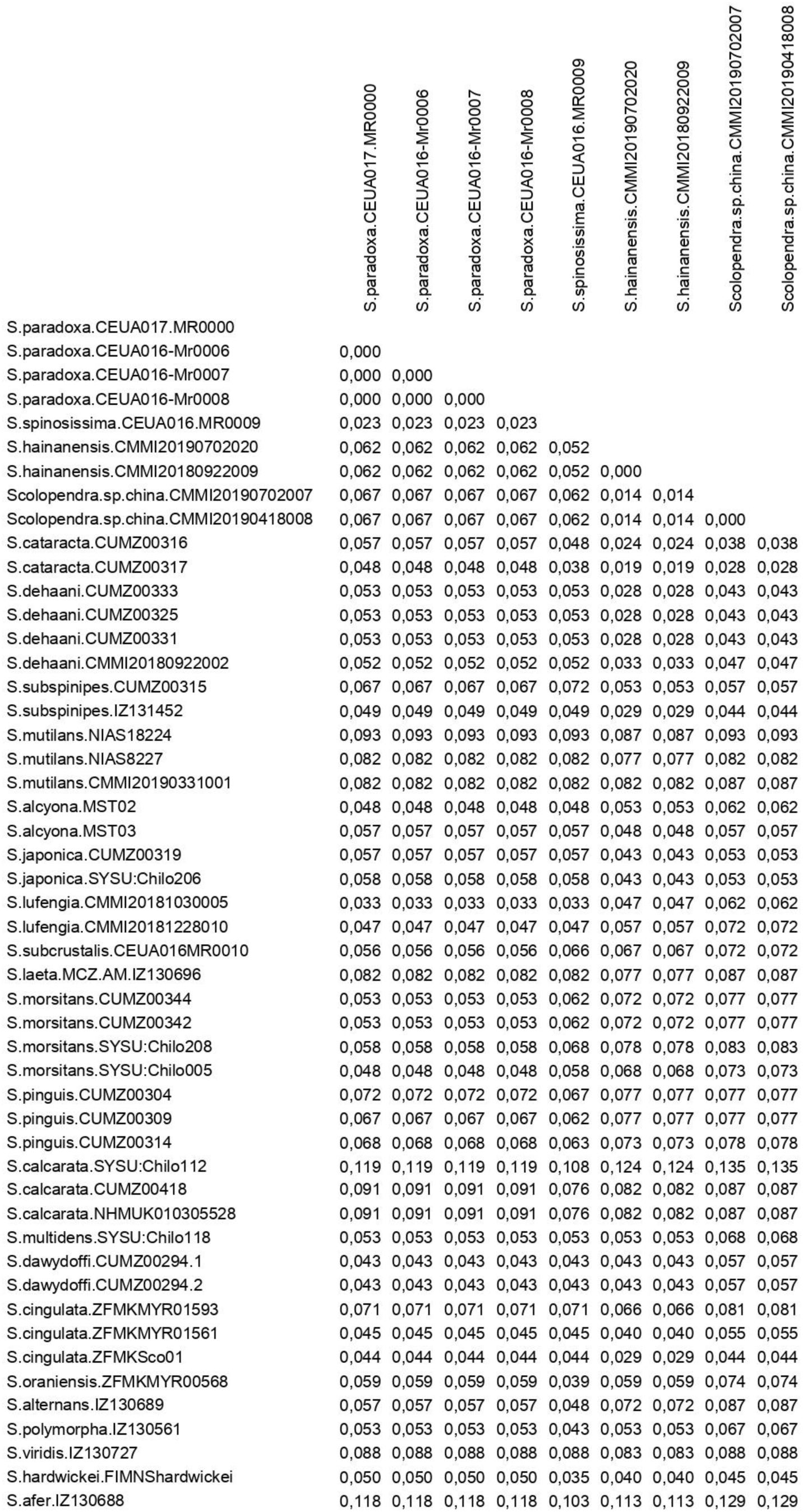
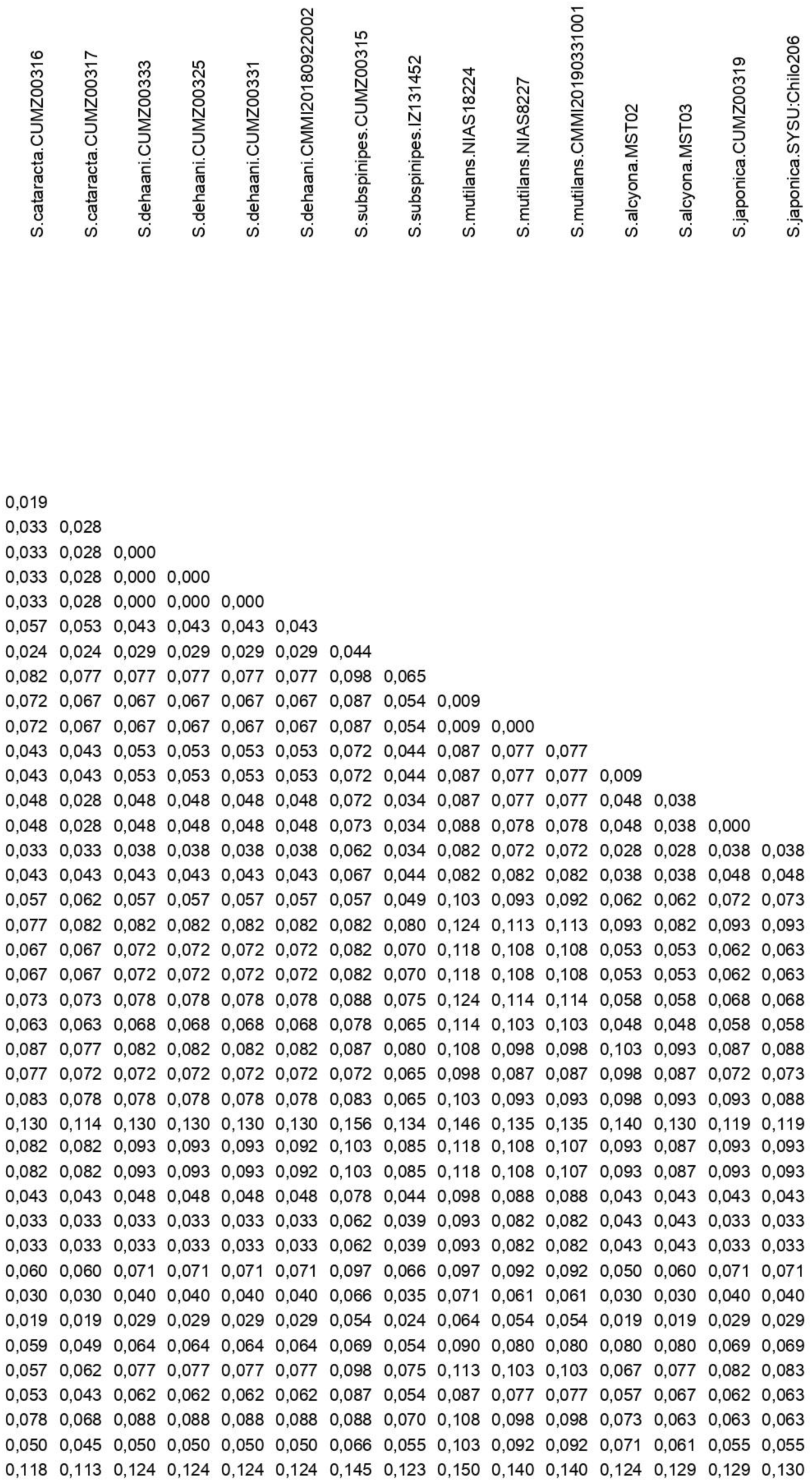
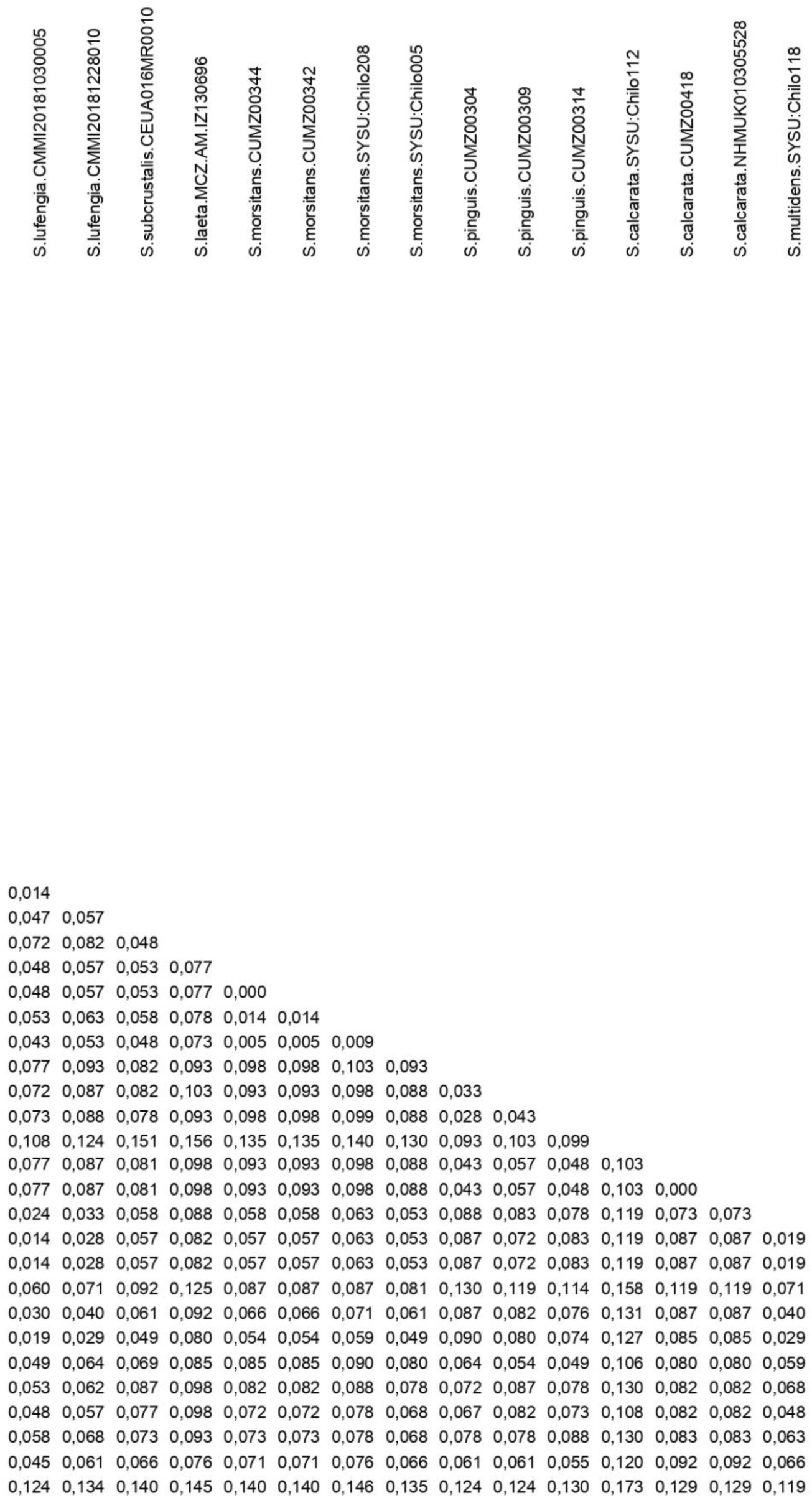
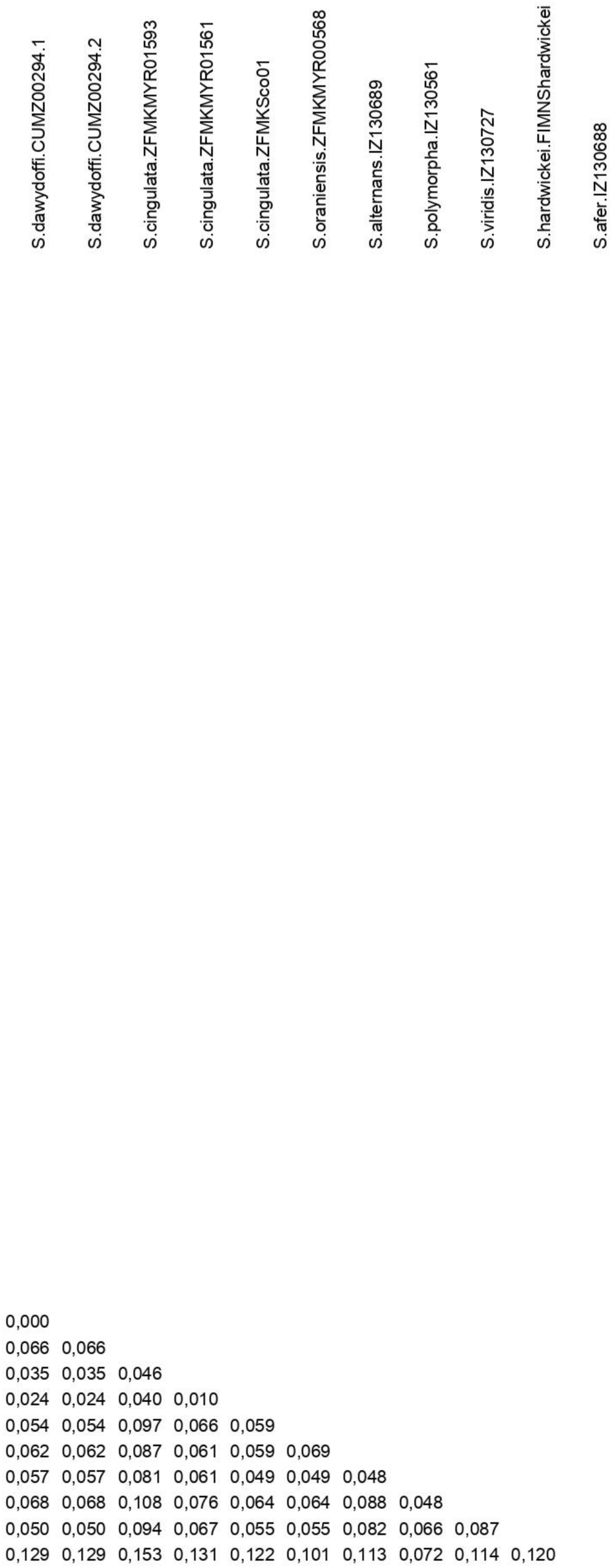
Divergence table obtained from 45 amino acid sequences of the partial COI gene belonging to 22 different species of the genus *Scolopendra*. Note that out of 820 alignments, only 14 exceptions to the five amino acids rule were found, eight of which correspond to sequences from the same species.

## References

Attems, C. (1930) Myriopoda. 2. Scolopendromorpha. In: Das Tierreich. Vol. 54. W. De Gruyter, Berlin and Leipzig, pp. 1–308.

Bonato, L., Chagas Junior, A., Edgecombe, G.D., Lewis, J.G.E., Minelli, A., Pereira, L.A., Shelley, R.M., Stoev, P. & Zapparoli, M. (2017) ChiloBase 2.0. A world catalogue of centipedes (Chilopoda). Disponible en: http://chilobase.biologia.unipd.it (Visitado el 19 Febrero 2024).

Bücherl, W. (1946) Novidades sistematicas na ordem Scolopendromorpha. Memorias do Instituto de Butantan, 19: 135–158.

Doménech, C. (2024) Type designation and redescription of *Scolopendra spinosissima* Kraepelin, 1903 (Scolopendromorpha, Scolopendridae), with remarks on related taxa. ZooKeys 1215: 311–334. 10.3897/zookeys.1215.129410

Doménech, C., Barbera, V.M. y Larriba, E. (2018) A phylogenetic approach to the Philippines endemic centipedes of the genus *Scolopendra* Linnaeus, 1758 (Scolopendromorpha, Scolopendridae), with the description of a new species. Zootaxa, 24; 4483(3): 401–427. 10.11646/zootaxa.4483.3.1. PMID: 30313775.

Edgecombe, G.D. & Giribet, G. (20087) Evolutionary biology of centipedes (Myriapoda: Chilopoda). Annual Review of Entomology, 27; 52:151–70. 10.1146/annurev.ento.52.110405.091326. PMID: 16872257.

Guindon, S., Dufayard, J.F., Lefort, V., Anisimova, M., Hordijk, W. y Gascuel, O. (2019) New algorithms and methods to estimate maximum-likelihood phylogenies: assessing the performance of PhyML 3.0. Systematic Biology, 59(3): 307–321. 10.1093/sysbio/syq010. PMID: 20525638.

Haase, E. (1887) Die Indisch-Australischen Myriopoden. 1. Chilopoden. Abhandlungen und Berichte des Königlichen zoologischen und anthropologisch-ethnographischen Museum zu Dresden, 4, 1–118.

Hall, T. (2011) BioEdit: An important software for molecular biology. GERF Bulletin of Biosciences, 2(1): 60–61.

Han, T., Lee, Y.B., Kim, S.H., Yoon, H.J., Park, I.G. y Park, H. (2018) Genetic variation of COI gene of the Korean medical centipede *Scolopendra mutilans* Koch, 1878 (Scolopendromorpha: Scolopendridae). Entomological Research, 12; 48(6): 559–566. 10.111/1748-5967.12331.

ICZN (1999) International Code of Zoological Nomenclature. Fourth Edition. The International Trust for Zoological Nomenclature, London, UK.

Kang, S., Liu, Y., Zeng, X., Deng, H., Luo, Y., Chen, K. & Chen, S. (2017) Taxonomy and identification of the genus *Scolopendra* in China using integrated methods of external morphology and molecular phylogenetics. Scientific Reports, 7, 16032. 10.1038/s41598-017-15242-7. PMID: 29167482; PMCID: PMC5700134.

Kimura, M. (1968) Evolutionary rate at the molecular level. Nature, 624–626. 10.1038/217624a0. ISSN: 0028-0836.

Kimura, M. (1983) The neutral theory of molecular evolution. Cambridge University Press; Cambridge, UK. 10.1017/CBO9780511623486

Koch, L. (1878) Japanesische Arachniden und Myriapoden. Beschrieben von. Dr. L. Koch in Nürnberg. Mit 2 Tafeln (XV u. XVI). Verhandlungen der kaiserlich-königlichen zoologisch-botanischen Gesellschaft in Wien; 27: 785–795.

Kraepelin, K. (1903) Revision der Scolopendriden. Mitteilungen aus dem Naturhistorischen Museum in Hamburg, 20(2): 1– 276.

Kronmüller, C. (2009) A new species of Scolopender from the Philippines (Chilopoda: Scolopendridae). Arthropoda, 17(1), 48–51.

Kronmüller, C. (2012) Review of the subspecies of Scolopendra subspinipes Leach, 1815 with the new description of the South Chinese member of the genus Scolopendra Linnaeus, 1758 named *Scolopendra hainanum* spec. nov. Spixiana, 35 (1) 19–27.

Kumar, S., Stecher, G., Li, M., Knyaz, C. y Tamura, K. (2018) MEGA X: Molecular evolutionary genetics analysis across computing platforms. Molecular Biology and Evolution, 1; 35(6): 1547–1549. 10.1093/molbev/msy096. PMID: 29722887; PMCID: PMC5967553.

Latreille, M. (1829) Les Myriapodes. Le régne animal distribué d’après son organisation, Nouvelle Édition, Revue et Augmentée. Tome IV. Crustacés, Arachnides et Partie des Insectes. Cuvier, PML, Paris, 4, 326–339.

Le, V.S., Dang, C.C. y Le, Q.S. (2017) Improved mitochondrial amino acid substitution models for metazoan evolutionary studies. BMC Evolutionary Biology, 12; 17(1):136. 10.1186/s12862-017-0987-y. PMID: 28606055; PMCID: PMC5469158.

Leach, W.E. (1816) A tabular view of the external characters of four classes of animals, which Linné arranged under Insecta; with the distribution of the genera composing three of these classes into orders, &c. and descriptions of several new genera and species. Transactions of the Linnean Society of London, 11, 306–400. 10.1111/j.1096-36642.1813.tb00065.x

Linnaeus, C. (1758) *Systema naturae per regna tria naturae, secundum classes, ordines, genera, species cum characteribus, diferentiis, synonymis, locis*. Tomus I. Editio Decima, Reformata. Salvii L, editor. Holmiae.

Lucas, H. (1840) Arachnides, Myriapodes et Thysanoures. Histoire Naturelle des Îles Canaries, 2(2): 49–52.

Lucas, H. (1853) Essai sur les animaux articulés qui habitent l’ile de Crète. Revue et Magasin de Zoologie Pure et Appliquée, 5(2): 418–424, 461-468, 514-531, 565-576.

Multiple Experiment Viewer [Internet]. [cited 18^th^ May 2024] Disponible en: www.mev.tm4.org.

National Center for Biotechnology Information (NCBI) [Internet]. [cited el 24^th^ January 2024] Disponible en: https://www.ncbi.nlm.nih.gov

Newport, G. (1844) A list of the species of Myriapoda order Chilopoda contained in the cabinets of the British Museum with synoptic descriptions of forty-seven new species. Annals and Magazine of Natural History, 13, 94–101. 10.1080/03745484409442576

Oeyen, J.P., Funke, S., Böhme, W. & Wesener, T. (2014) The evolutionary history of the rediscovered Austrian population of the giant centipede *Scolopendra cingulata* Latreille 1829 (Chilopoda, Scolopendromorpha). PLoS ONE, 9(9), e108650. 10.1371/journal.pone.0108650

Pocock, R.I. (1891). On the Myriopoda of Burma. Part 2. Report upon the Chilopoda collected by Sig. L. Fea and Mr. E. W. Oates. Annali del Museo Civico di Storia Naturale di Genova, (ser.2)10: 401–432.

Porat, C. (1876) Om några exotiska Myriopoder. Bihang till Kongliga Svenska Vetenskaps-Akademien Handligar, 4, 1–48.

Ratnasingham, S. y Hebert, P.D.N. (2007) BOLD: The Barcode of Life Data System (www.barcodinglife.org). Molecular Ecology Notes, 24; 7: 355–364. 10.1111/j.1471-8286.2006.01678.x.

Rice, P., Longden, I. y Bleasby, A. (2000) EMBOSS: The European Molecular Biology Open Software Suite. Trends Genetics, 16(6):276–277. 10.1016/s0168-9525(00)02024-2

Rogers, H.H. y Griffiths-Jones, S. (2012) Mitochondrial pseudogenes in the nuclear genomes of *Drosophila*. PLoS One, 7(3):e32593. 10.1371/journal.pone.0032593. PMID: 22412894; PMCID: PMC3296715.

Say, T. (1821). Description of the Myriapoda of the United States. Journal of the Academy of Natural Sciences of Philadelphia, 2 (1): 102–114

Schileyko, A.A., Vahtera, V. & Edgecombe, G.D. (2020) An overview of the extant genera and subgenera of the order Scolopendromorpha (Chilopoda): a new identification key and updated diagnoses. Zootaxa, 4825(1), 001–064. 10.11646/zootaxa.4825.1.1

Siriwut, W., Edgecombe, G.D., Sutcharit, C. & Panha, S. (2015) The centipede genus *Scolopendra* in mainland Southeast Asia: molecular phylogenetics, geometric morphometrics and external morphology as tools for species delimitation. PLoS ONE, 10 (8), e0135355. 10.1371/journal.pone.0135355. Erratum in: *PLoS One*. 2015; 10(9):e0139182. PMID: 26270342; PMCID: PMC4536039.

Siriwut, W., Edgecombe, G.D., Sutcharit, C., Tongkerd, P. & Panha, S. (2016) A taxonomic review of the centipede genus *Scolopendra* Linnaeus, 1758 (Scolopendromorpha, Scolopendridae) in mainland Southeast Asia, with description of a new species from Laos. ZooKeys, 590, 1–124. 10.3897/zookeys.590.7950 PMID: 27408540; PMCID: PMC4926625.

Susko, E. y Roger, A.J. (2021) Long Branch Attraction Biases in Phylogenetics. Systematic Biology, 70(4): 838–843. 10.1093/sysbio/syab001

Undheim, E.A. y King, G.F. (2011) On the venom system of centipedes (Chilopoda), a neglected group of venomous animals. Toxicon, 15;57(4):512–24. doi: 10.1016/j.toxicon.2011.01.004.

Undheim, E.A., Fry, B.G. y King, G.F. (2015) Centipede venom: recent discoveries and current state of knowledge. Toxins, 25;7(3):679–704. 10.3390/toxins7030679. PMID: 25723324; PMCID: PMC4379518.

Vahtera, V. y Edgecombe, G.D. (2014) First molecular data and the phylogenetic position of the millipede-Like centipede *Edentistoma octosulcatum* Tömösváry, 1882 (Chilopoda: Scolopendromorpha: Scolopendridae). PLoS ONE, 9(11): e112461. 10.1371/journal.pone.0112461.

Vahtera, V., Edgecombe, G.D. & Giribet, G. (2013) Phylogenetics of scolopendromorph centipedes: can denser taxon sampling improve an artificial classification*?* Invertebrate Systematics, 27 (5), 578–602. 10.1071/IS13035.

Vahtera, V., Edgecombe, G.D. y Giribet, G. (2012) Evolution of blindness in scolopendromorph centipedes (Chilopoda: Scolopendromorpha): insight from an expanded sampling of molecular data. Cladistics, 28, 4–20.

Waldock, J. y Edgecombe, G.D. (2012) A new genus of scolopendrid centipede (Chilopoda: Scolopendromorpha: Scolopendrini) from the central Australian deserts. Zootaxa, 13; 3321: 22–36. 10.5281/zenodo.281187

Weblogo [Internet]. [citado el 17 de Mayo 2024] Disponible en: https://weblogo.berkeley.edu/logo.cgi.

Wood, H.C. Jr. (1861) Description of new species of *Scolopendra* in the collection of the Academy. Proceedings of the Academy of Natural Sciences of Philadelphia, 10–15.

Xia, X. (2009) *Assessing substitution saturation with DAMBE*. In: The phylogenetic handbook: a practical approach to phylogenetic analysis and hypothesis testing, Philippe Lemey, Marco Salemi, and Anne-Mieke Vandamme (eds.) Published by Cambridge University Press. UK.

Zhang, C.Z. y Wang, K-Q. (1999) A new centipede *Scolopendra negrocapitis* sp. nov. from Hubei Province, China (Chilopoda: Scolopendromorpha: Scolopendridae). Acta Zootaxonomica Sinica, 24: 136–137. 10.3969/j.issn.1000-0739.1999.02.003

Zuckerkandl, E. y Pauling, L.B. (1962) *Molecular disease, evolution, and genic heterogeneity*. In: Kasha, M. and Pullman, B., Eds., Horizons in Biochemistry, Academic Press, New York, USA,189–225.

